# Time and Space in Segmentation

**DOI:** 10.1101/2020.09.12.294611

**Authors:** Erik Clark

## Abstract

Arthropod segmentation and vertebrate somitogenesis are leading fields in the experimental and theoretical interrogation of developmental patterning. However, despite the sophistication of current research, basic conceptual issues remain unresolved. These include (1) the mechanistic origins of spatial organisation within the segment addition zone (SAZ); (2) the mechanistic origins of segment polarisation; (3) the mechanistic origins of axial variation; and (4) the evolutionary origins of simultaneous patterning. Here, I explore these problems using coarse-grained models of cross-regulating dynamical processes. In the morphogenetic framework of a row of cells undergoing axial elongation, I simulate interactions between an “oscillator”, a “switch”, and up to three “timers”, successfully reproducing essential patterning behaviours of segmenting systems. By comparing the output of these largely cell-autonomous models to variants that incorporate positional information, I find that scaling relationships, wave patterns, and patterning dynamics all depend on whether the SAZ is regulated by temporal or spatial information. I also identify three mechanisms for polarising oscillator output, all of which functionally implicate the oscillator frequency profile. Finally, I demonstrate significant dynamical and regulatory continuity between sequential and simultaneous modes of segmentation. I discuss these results in the context of the experimental literature.

## 1 Introduction

Arthropod segmentation [1] and vertebrate somitogenesis [2, 3] are paradigmatic examples of developmental pattern formation. They involve the coordination of diverse cellular processes – signalling, gene expression, morphogenesis – yet can be conveniently abstracted to a single dimension, making them conducive to mathematical modelling and theoretical analysis.

The spatial patterns produced by segmentation processes have at least three components (Figure 1A). First, *periodicity* – the reiterated nature of individual metameres. Second, the intrinsic *polarity* of these units, evident in the expression of segment-polarity genes. Finally, *regionalisation* – variation in the length and identity of metameres according to their position along the anteroposterior (AP) axis. Segmentation patterns are highly robust within species [4–6] but evolutionarily flexible between species [7–10], with differences in the number, size, and specialisation of segments contributing to the considerable morphological diversity of segmented clades.

**Figure 1:**
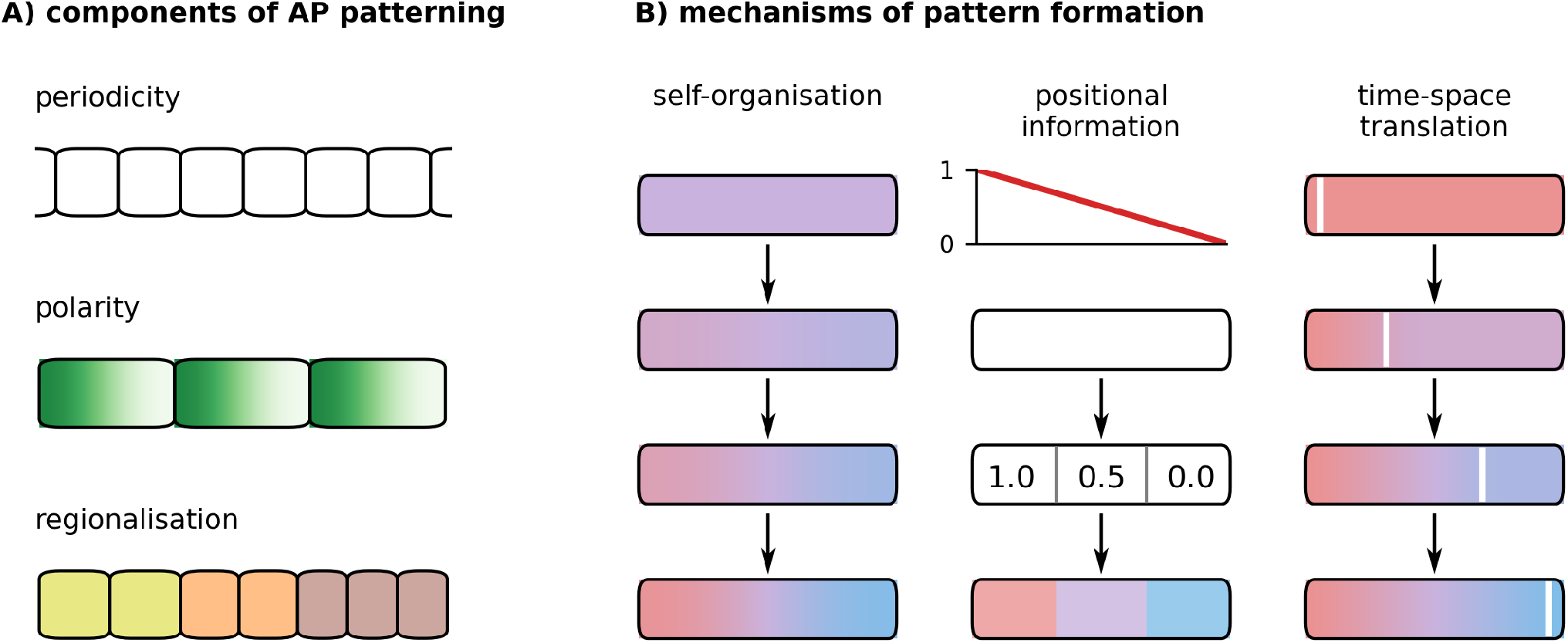
Key patterning concepts. **A**: Three components of AP patterning: periodicity (metamerism); intra-segment polarisation; region-specific segment properties (size, gene expression). **B**: Three mechanisms for spatial pattern formation. Left, organisation emerges in an initially homogenous tissue by local reaction and diffusion. Middle: an embryonic field (white) interprets positional information (red line) to acquire positional values, and subsequently differentiates (coloured domains). Right, a wavefront (white line) moves across a tissue as it matures from an early state (red) to a late state (blue), and freezes the temporal pattern in space.

Segment patterning emerges, in most species, from fundamentally temporal developmental mechanisms. In both vertebrates and arthropods, temporal periodicity is generated by a “segmentation clock” (molecular oscillator), then translated into spatial periodicity by a “wavefront” of cell maturation that sweeps the elongating AP axis from anterior to posterior at a tightly controlled rate [11–17]. These tissue-scale dynamics are generated, at the molecular level, by the characteristic rates, time delays, and flow properties of intercellular and intracellular regulatory networks [18–23], in combination with morphogenetic processes [24–26].

Genetic, environmental, and pharmacological perturbations can alter the segment pattern [13, 27–37], providing insight into its mechanistic basis. However, despite the experimental sophistication of vertebrate research in particular, these approaches are often constrained by functional redundancy and pleiotropic effects. Modelling studies provide a complementary way to interrogate segmentation processes, and have produced important conceptual advances [11, 18, 38, 39]. Yet a range of modelling approaches with contrasting regulatory bases have been able to give more or less convincing approximations of empirical observations, and it can be unclear how to best choose between them [39–50].

In this context, it is a worthwhile exercise to step away from the particularities of specific genes, signals, and experimental species, and consider the generic ramifications of basic assumptions. Here, I identify four aspects of the segmentation process where the nature of patterning is open to question or unclear. I then explore each of these issues using coarse-grained, illustrative models, in which potentially complex processes are represented by interactions between phenomenological “dynamical modules”. Finally, I discuss the implications of these results in the context of the experimental literature.

## 2 Patterning problems in segmentation

### 2.1 Sources of spatial information

Developmental patterning requires that cells’ state trajectories (changes in internal state over time) are influenced by their location – either their global position within the tissue or embryo, or their local position relative to certain other cells. In other words, some cell state variable must respond to a source of spatial information, and different values for this variable must cause otherwise identical cells to diverge in fate. Three general mechanisms that accomplish this have been identified:

1. **Self-organisation** (i.e. reaction-diffusion) mechanisms [51–53] (Figure 1B, left), in which small heterogeneities between cells, perhaps originating ultimately from stochastic fluctuations, are amplified by coupled positive and negative feedback loops into regular spatial patterns with a characteristic length scale. The key here is that the feedback loops and diffusive processes effect a local exchange of state information between cells, so that state trajectories become dependent on relative position.
2. **Positional information** (PI) mechanisms [54–56] (Figure 1B, middle), in which a polarised scalar variable (*e*.*g*., the concentration of a graded morphogen) provides a coordinate system for a bounded embryonic field. Cells “interpret” the positional information to acquire a “positional value”; the value serves as a proxy for a cell’s position within the field, and influences its subsequent state trajectory. PI processes are constrained by the accuracy and precision with which the PI signal is established and interpreted.
3. **Time-space-translation** (TST) mechanisms [11, 57–61] (Figure 1B, right), in which a tissue-scale velocity converts a temporal pattern into an isomorphic pattern arrayed across space. The velocity (*e*.*g*., a wave of gene expression across a field of cells, or a flow of cells relative to the source of an extracellular signal) causes cells at each position to experience a particular input at a different time relative to ongoing changes to cell state, with correspondingly different effects on cell fate.

These modes of patterning are conceptually distinct, but are unlikely to be cleanly dissociable in real systems. Cell states change constantly as the result of intrinsic processes (*e*.*g*., the synthesis and turnover of gene products) and extrinsic influences (*e*.*g*., the transduction of extracellular chemical and mechanical signals), and, in an embryonic context, thousands of cells respond to each other’s signals and move relative to one another simultaneously. The massively parallel feedback loops between cell state, signalling and morphogenesis, occurring over a wide range of timescales and lengthscales [62], mean that non-trivial developmental phenomena are unlikely to be governed by a single patterning mechanism, but will instead emerge from a recursive blend of self-organisation, PI, and TST. The challenge of developmental biology is to unpick these processes and render the whole system comprehensible. Indeed, perhaps it is in the interactions *between* patterning mechanisms that explanations for developmental robustness and evolutionary flexibility are to be found [63].

### 2.2 SAZ organisation and the “determination front”

With some significant exceptions (see below), segmentation occurs in the context of posterior elongation, with segments/somites emerging sequentially from the anterior of a segment addition zone (SAZ, arthropods) or the presomitic mesoderm (PSM, vertebrates). [Unless otherwise indicated, “SAZ” is hereafter used as a generic term that encompasses both SAZ and PSM.]

In both arthropods and vertebrates, the posterior tip of the SAZ continuously moves away from more anterior structures, either by a mix of convergence extension movements and cell proliferation (arthropods) [64–68], or by cell influx, cell proliferation and a greater motility of posterior *vs* anterior cells (vertebrates) [26, 69–74]. These processes, and therefore the maintenance of axis elongation, depend directly or indirectly on a posterior signalling centre (PSC) involving Wnt signalling, itself maintained by positive feedback loops [32, 75–85]. The posterior signalling environment maintains SAZ cells in an immature, unpatterned state, preventing their differentiation into segmental fates [16, 30, 32, 86–89].

The more anterior the location of a cell within the SAZ, the more remote it is from the PSC, both in terms of physical distance, and in terms of time. (Note that the SAZ is a dynamic structure through which there is a continuous flux of cells.) At a certain point within the SAZ, cells undergo a significant state transition, and begin expressing different sets of transcription factors (*e*.*g*., Odd-paired rather than Dichaete/Sox21b and Caudal in arthropods [16, 17, 85, 90–94]; Paraxis/TCF15 *vs* Mesogenin, and a combinatorial code of T-box factors in vertebrates [31, 95– 100]). The transition coincides with the specification of segmental fate, at least partially because the transcription factors on either side of it differentially regulate segmentation gene enhancers [98, 101–106]. This transition or “determination front” [13] can therefore be identified with the wavefront of the clock and wave-front model.

The positioning of the determination front – and the origins of spatial organisation within the SAZ more generally – is fundamental to the control of segmentation. It is clear that SAZ geometry is affected by the levels of Wnt and other signals within the embryo [3, 107]. Less clear, however, are the respective roles played by spatial and temporal information in producing these outcomes. The more anterior a cell is, the further it is from the PSC and the closer it is to patterned tissue that may produce antagonistic signals such as retinoic acid [108]. However, it will also have spent a longer time degrading posterior signals [30, 32, 109] and potentially undergoing other autonomous changes in cell state. Because temporal and spatial coordinate systems are overlaid on one another in this way, they are extremely difficult to disentangle.

There are two distinct regulatory questions here. The first, and simplest, asks to what extent the signalling environment along the SAZ is shaped by position dependent, diffusive effects, as opposed to advection (bulk transport) and temporal decay [46]. While both play some role, the latter effects appear to dominate in embryos. For example, Wnt perturbations take hours to affect somitogenesis in zebrafish, with effects on SAZ gene expression travelling from posterior to anterior at the rate of cell flow [88], and dissected PSM tissue generally maintains an endogenous maturation and patterning schedule [110–113]. In contrast, most segmentation models assume that SAZ patterning is governed by a characteristic lengthscale, rather than by a characteristic timescale.

The second question, less frequently raised, concerns the relationship between the signalling levels a cell experiences, and the dynamics of its internal state. Most segmentation models implicitly assume a PI framework, in which a cell moves within a PI field, continuously monitoring a graded signal, until a particular positional value induces a change in cell state [39, 41, 47, 49, 114, 115]. (Other models propose that cells actually respond to a different property of the field [44, 48], but the basic principle of decoding a PI signal remains the same.) An alternative hypothesis is that cells travel autonomously along a maturation trajectory from a “posterior” state to an “anterior” state over a period of time, with the signalling environment governing the rate of cell state change, but not determining cell state *per se*.

The two hypotheses predict quite different behaviour: the former suggests that intermediate cell states are stable in intermediate signalling environments and that cell maturation is associated with the decoding of a specific external trigger; the latter suggests that intermediate cell states are inherently unstable and that signalling levels and cell state can be partially decoupled. Significantly, recent results from stem cells and *ex vivo* cultures support the latter hypothesis by pointing to an important role for cell-intrinsic dynamics: stems cells in constant culture conditions will transition from a posterior PSM-like state to an anterior PSM-like state over a characteristic period of time [100, 116, 117], while more anterior PSM cells will oscillate slower than posterior PSM cells in otherwise identical culture conditions [118].

Thus, the emphasis in the modelling literature on SAZ lengthscale and morphogen decoding is at odds with empirical findings. However, because much of axial elongation can be approximated by a pseudo-steady state scenario [39] in which the dimensions of the SAZ remain constant and spatial and temporal coordinate systems coincide (and are therefore accurate proxies for one another), timescale-based and lengthscale-based models often yield similar predictions. A central aim of this study is to explicity contrast these two patterning frameworks and determine their characteristic implications.

### 2.3 Prepatterns and polarity

Segments and somites both have an inherent polarity, but their polarity manifests in different ways. In arthropods, the segmentation process generates a segment-polarity repeat with three distinct expression states (*i*.*e*., anterior, posterior, and a buffer state in between), with polarised signalling centres forming wherever the anterior and posterior states abut [119–123]. These signalling centres demarcate the boundaries of the future (para)segments, and later pattern the intervening tissue fields by a PI mechanism [124–127].

In vertebrates, the segment-polarity pattern is just a simple alternation of anterior and posterior fates [128, 129], and is not necessary for boundary formation *per se* [130–132]. Instead, somitic fissure formation depends on the relative timing of cells’ mesenchymal to epithelial transition (MET), and differences in their functional levels of specific extra-cellular matrix (ECM) components such as Cadherin 2 (CDH2) [133]. Both are modulated by the segmentation clock [134, 135], explaining the regularity, scaling properties, and species-specificity of somite size, as well as the tight coordination between somite boundary formation and the segment polarity pattern. However, note that there is spatial information in the ECM pattern that is absent from the segment-polarity pattern [136] — this explains why the somitic fissures that gate cohorts of cells into coherent morphological structures form only at P-A segment-polarity interfaces, and not also at the A-P interfaces in between, as would be predicted by a prepattern mechanism.

Thus, arthropod segments are polarised because their morphogenesis is specified by a polarised prepattern, while vertebrate somites are polarised because morphogenesis intersects with a non-polarised prepattern in a polarised way. However, in both cases the origin of this polarity still needs to be explained. In all vertebrates and at least some arthropods, the segmentation clock is thought to be based around coupled oscillations of *her* /*hes* gene expression and Notch signalling [28, 29, 38, 89, 137–147]; both theoretical models and empirical data suggest that such networks generate sinusoidal oscillations, which are not inherently polarised [18, 44, 89, 116, 142, 148, 149]. The problem thus arises of how a non-polarised pattern in time is able to specify a polarised pattern in space. I will present two strategies for how this could be achieved, one inspired by data from arthropods, and the other by data from vertebrates.

### 2.4 Axial variation and scaling

In real embryos, elongation, SAZ maintenance and segmentation eventually terminate, usually all at roughly the same time, after the production of a species-specific and relatively invariant number of segments [97, 150]. These segments often have different sizes and molecular identities according to their axial position; for example, tail segments are usually smaller than trunk segments and express more 5’ Hox genes. Understanding how segmentation “profiles” are generated requires determining (1) which parameters of the segmentation process are modulated to produce axial variation and terminate segment addition, and (2) how the dynamics of these parameters are controlled.

#### Segment identity and segmentation duration

In both vertebrates and arthropods, segment identities are specified by sequentially expressed Hox genes [151–155]; in at least some arthropods, Hox gene expression is regulated by another set of sequentially expressed transcription factors, encoded by the gap genes [156, 157]. Hox and gap gene expression is established in parallel with, but independent of, segment boundary patterning. Indeed, segment number and segment identity can be decoupled by various perturbations [21, 34, 158–160].

Hox and gap gene expression is activated by posterior factors such as Cdx and Wnt [16, 161–164], and the individual genes also cross-regulate each other’s expression. Thus Hox and gap gene expression depends partly on the state of the SAZ and partly on intrinsic dynamics [58, 165–167]. As well as generating regionalisation information, the sequential expression of these genes affects segment patterning, via effects on SAZ maintenance and axial elongation [78, 168–171].

Reaction-diffusion processes [46, 47, 150] could additionally play a role in controlling the duration of segmentation, as could the proliferation rate and initial size of unpatterned tissues [72, 172]. Thus, segmentation termination is likely controlled by some kind of effective timer, but it is not clear whether this timer resides in the dynamical properties of a particular gene network, or is broadly distributed in the system at large.

#### Segment size control

In a clock and wavefront framework, segment length is determined jointly by clock period and wavefront velocity [11, 39]. Both parameters change over the course of embryogenesis [87, 97, 150, 173], but the change in oscillation period is opposite of that expected from changes in segment size, increasing at late stages while segment sizes decrease. Thus, variation in the clock is more important for explaining segment size variation between species [23, 97, 174, 175] than the size variation in an individual embryo over time. In addition, the oscillation period is unaffected by embryo size variation, although such variation has a strong effect on segment length [47]. Therefore, variation in segment size over time and between individuals must be largely a consequence of variation in wavefront velocity.

Wavefront velocity is generally similar to – but not necessarily identical with – the axial elongation rate; discrepancies between the two velocities reflect temporal variation in SAZ length [149]. Explaining how the segment profile is generated therefore requires understanding how elongation rate and SAZ length are regulated over time. Are changes to these properties linked to Hox gene expression changes or independent of them? Are elongation rate and SAZ length controlled separately or are they mechanistically linked? What accounts for observed scaling relationships between segment length, SAZ length, and oscillator wave patterns [5, 44, 47, 97, 149, 173]? Within species, these scaling relationships are broadly (though not entirely [176, 177]) conserved between individuals, despite variation in embryo size and strong temperature-dependence of developmental rates. At the same time, they vary moderately within individuals over the course of segmentation, and can vary dramatically across species.

Determining the upstream temporal regulation and proximate mechanistic causes of axial variation is central to questions of developmental robustness and evolutionary flexibility. At the same time, by providing a more exacting set of observations for segmentation models to explain, the investigation of these issues should also generate considerable insight into the control of segmentation at “steady state”. I will demonstrate this point by showing that the downstream effects of temporal variation in elongation rate and SAZ size depend on whether the SAZ is patterned by spatial or temporal information.

### 2.5 Simultaneous segmentation and its evolution

Finally, in several groups of holometabolous insects (including the best studied segmentation model, *Drosophila*) the ancestral segmentation process has been modified by evolution to such an extent that it is almost unrecognisable [178]. The segmentation clock has been replaced by stripe-specific patterning by gap genes [179–184], while segments mature near-simultaneously at an early stage of embryogenesis, rather than sequentially over an extended period of time. Contrasting with the temporal nature of clock and wavefront patterning, the *Drosophila* blastoderm has become one of the flagship models of positional information [185–187].

Yet, in recent years it has been found that many of the dynamical aspects of *Drosophila* segmentation gene expression are strikingly reminiscent of sequential segmentation. For example, domains of gap gene and pair-rule gene expression move from posterior to anterior across nuclei over time [123, 188–192], while the timing of pattern maturation is regulated by factors associated with the SAZ in sequentially-segmenting arthropods [17, 106, 193]. At the same time, there have been several experimental results that belie a strict PI framework, such as those characterising the activation thresholds of Bicoid target genes [194, 195], or the information content of maternal gradients [185, 186, 196].

Clarifying the regulatory homologies between simultaneous and sequential segmentation and determining the functional significance of *Drosophila* segmentation gene dynamics will shed light on both simultaneous and sequential segmentation processes, as well as the evolutionary trajectories between them. I will delineate the regulatory changes and initial conditions required to maintain segment patterning in the absence of elongation, providing an initial framework for such efforts.

## 3 Modelling framework

The four issues described above – (1) the origins of spatiotemporal order within the SAZ, (2) the origins of segment polarisation, (3) the proximate and ultimate regulatory causes of axial variation, and (4) the existence of extreme transmogrifications of the segmentation process – are as much questions about the origins and transformations of spatial information as they are questions about particular biological mechanisms. To focus attention on spatial information, I have chosen to use a modelling framework that simulates the dynamics and interactions of regulatory processes within cells but remains agnostic as to their molecular implementation. To easily follow the logic of patterning, the models are simple, uni-dimensional, and deterministic.

Unlike models that explicitly specify the properties of the patterning field, here the emphasis is on internal cell state and on autonomously-driven cellular dynamics. An “embryo” is modelled as a 1D array of largely autonomous “cells”; the state of each cell is represented by a vector of state variables, each of which represents a particular “dynamical module” (Figure 2A). For simulations, I use discrete time and synchronous update: the state of a cell at *t*_*n*+1_ is calculated from its state at *t*_*n*_ according to pre-specified regulatory rules, which are identical for all cells.

**Figure 2:**
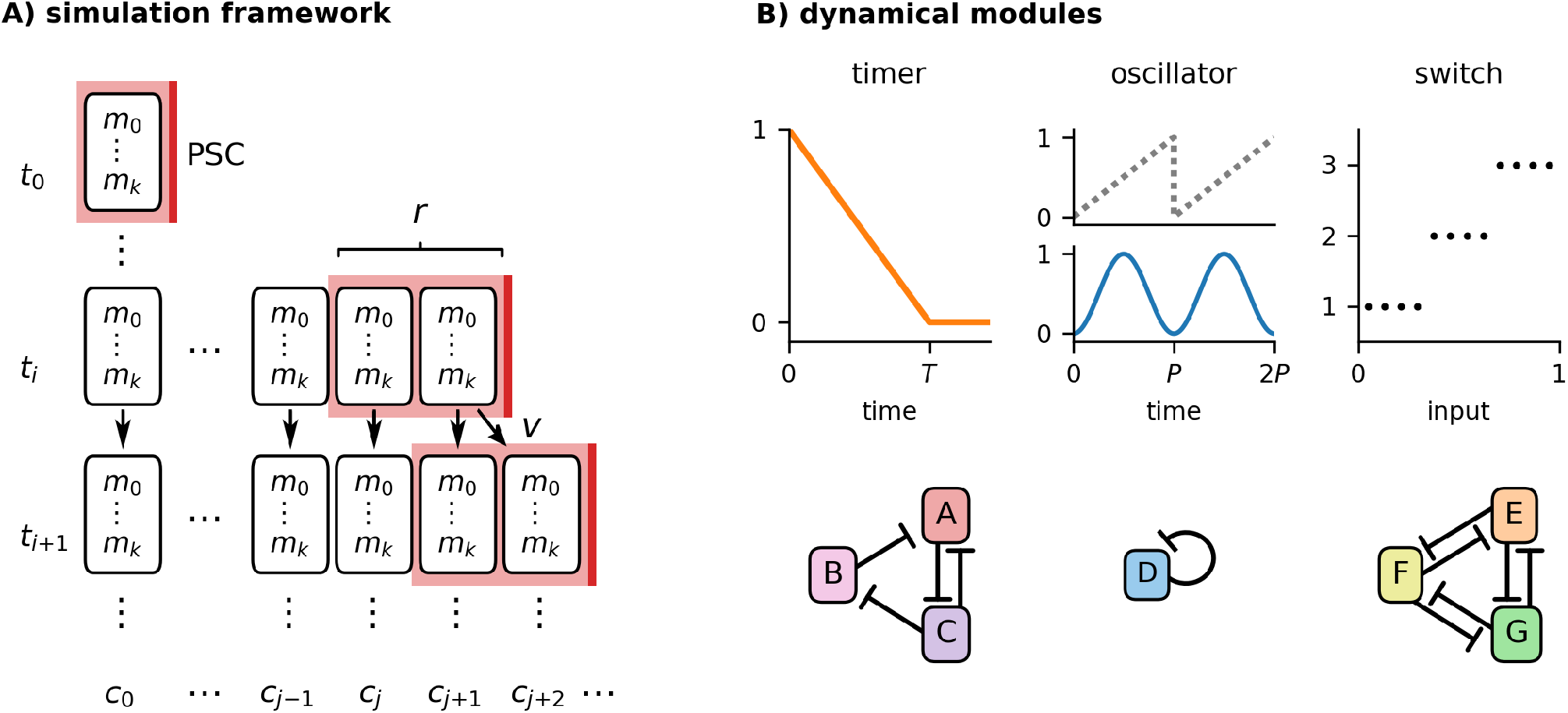
Modelling framework. **A:** Diagram of a model simulation. Each row (*t*_0_, *t*_1_, …) is a timepoint; each column (*c*_0_, *c*_1_, …) is a cell. Each cell is characterised by a vector of dynamical modules (*m*_0_, …, *m*_*k*_) whose states are at updated each timepoint. Posterior to the last cell is a posterior signalling centre (“PSC”, red lines) with a spatial range (pink domain) of *r* cells. The embryo starts at *t*_0_ with a length of one cell, then grows at rate *v* by duplicating its most posterior cell. **B:** The three types of “dynamical modules” used in the models. A *timer* has a duration *T*. An *oscillator* has a hidden phase (grey dotted line), an output level (blue line) and a period, *P*. A *switch* has *n* mutually exclusive states (here *n* = 3), each activated by different input conditions. Module behaviours are assumed to derive from underlying cellular regulatory networks, including (for example) AC-DC motifs [197] (left), autorepression (middle), or mutual repression (right).

Each dynamical module represents a set of cellular components, such as a cross-regulatory gene network, that executes a coherent dynamical behaviour. The modules may be of three general types. A “timer”, *τ*, (Figure 2B, left) decreases from an initial state 1 to a final state 0 at rate 1*/T* (where *T* is the timer duration in timesteps). [*N*.*B*., this is just an abstract representation of a canalised cell state trajectory and should not be confused with linear decay of a particular gene.] An “oscillator”, *o*, (Figure 2B, middle) generates a sinusoidal output: it has a hidden phase value *ϕ* [0, 1] which advances at rate 1*/P* (where *P* is the period in timesteps), and is converted to a functional output level by the function *o* = (sin(2*π*(*ϕ* −1*/*4)) + 1)*/*2. Finally, a “switch”, *s*, (Figure 2C, right) remains at an initial state 0 until one of *n* mutually exclusive states is activated by a rule. Each module may additionally be regulated by the current state of one or more different modules, allowing for flexible context-specific behaviour.

Each embryo has an initial length *L*_0_ (usually 1 cell) and a polarity defined by a posterior signalling centre (PSC) that is extrinsic to the row of cells. The PSC emits a Boolean posterior signal that has a spatial range of *r* cell diameters. The signal can affect cell state by entering as a variable in the regulation of a dynamical module. It also induces embryonic elongation. Elongation is effected within the model by duplicating the posterior-most cell (including all its state variables) at regular time intervals determined by an elongation rate *v* (*e*.*g*., if *v* = 0.2 the embryo adds a cell every 5 timesteps). Each increase in embryonic length posteriorly displaces the PSC by one cell diameter, freeing the (*L − r*)_*th*_ cell from its influence. (This elongation mechanism is simply a modelling convenience, representing any combination of cell proliferation, cell rearrangement, or cell recruitment.)

For many of the simulations, no other morphogenetic or cell communication processes are included, and initial conditions are identical between cells. As a consequence, the elongation-driven dynamics of the posterior signal (*i*.*e*., a velocity) is the sole source of spatial information in the system; all patterning must occur via TST (time-space translation). In other simulations, cells are allowed to calculate their spatial distance from the posterior signal or to have different initial conditions, introducing PI effects. Self-organisation effects are excluded from the models so as to more clearly contrast PI and TST, but they are undoubtedly important in real segmenting tissues, particularly for counteracting stochasticity.

The series of models that I investigate is summarised in Supplementary Figure 1. Mathematical details for each model and a list of the parameter values used in each simulation are provided in a Supplementary Document. Simulations were implemented in Python, using the libraries NumPy [198] and Matplotlib [199].

## 4 A clock and timer model

The first model (Figure 3A) presents a simple clock and wavefront scenario, which will be iterated on in later sections. Here, the main concern is establishing the basic logic of the model and understanding the effect of each parameter on model behaviour. I contrast two variants of the model, one in which the SAZ is patterned by temporal information, and another in which the SAZ is patterned by spatial (positional) information. Their patterning output is overall similar, but they are differently affected by the elongation rate.

**Table 1:** Table of model symbols.

| Symbol | Meaning |
| --- | --- |
| $t$ | time |
| $x$ | AP position |
| $\tau_{(1)}$ | timer (1) |
| $T_{(1)}$ | duration of timer (1) |
| $\lambda$ | SAZ lengthscale |
| $o$ | oscillator level |
| $\phi$ | oscillator phase |
| $P$ | oscillator period |
| $s$ | switch |
| $n$ | number of switch states |
| $L$ | embryo length |
| $L_{\text{SAZ}}$ | SAZ length |
| $L_{\text{seg}}$ | segment length |
| $v$ | elongation rate |
| $r$ | signal range |
| $\theta$ | oscillator threshold |
| $k$ | feedback shape parameter |

**Figure 3:**
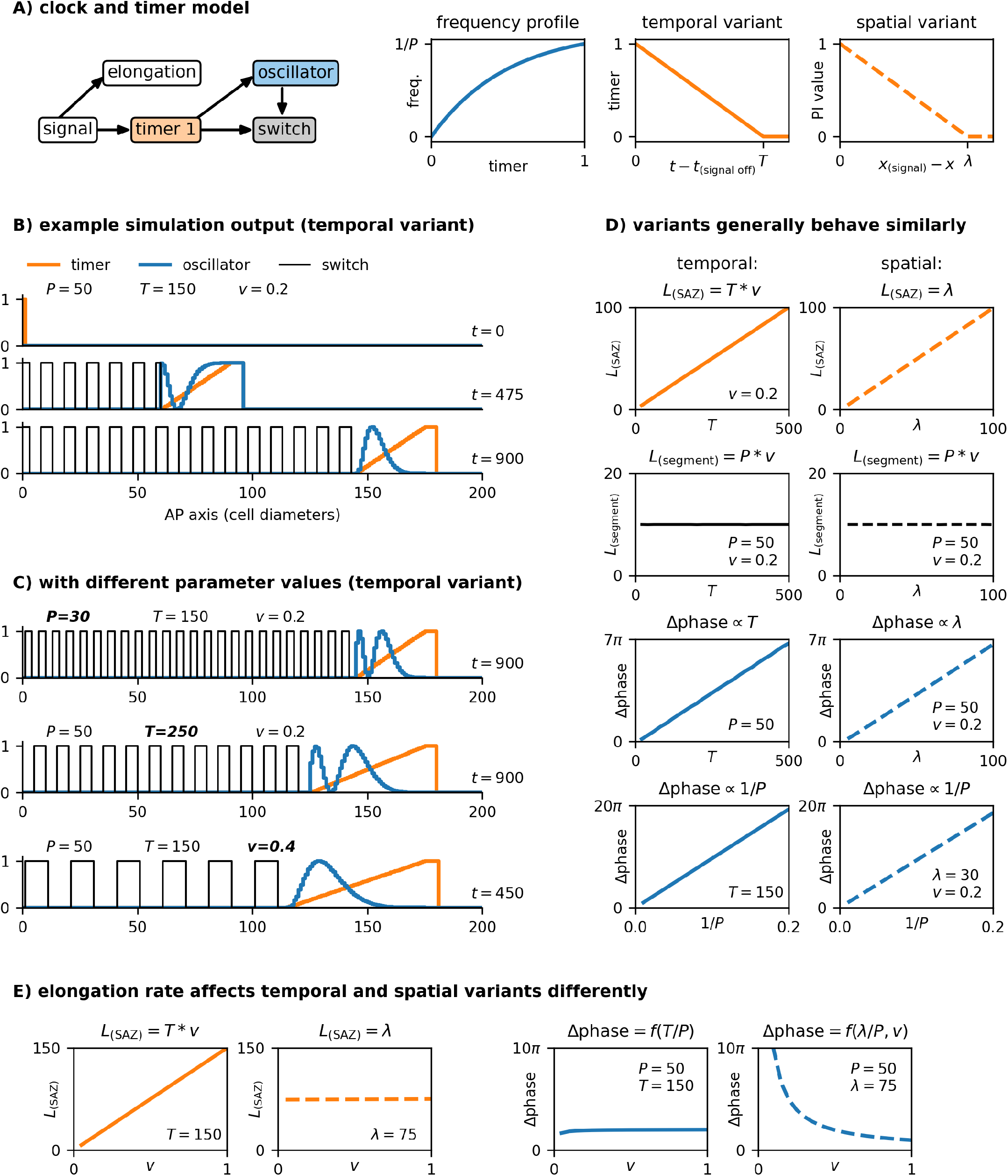
Clock and timer model. **A:** Summary of model. From left to right: diagram of model structure (dynamical modules shown as coloured boxes, other model features shown as white boxes, regulatory connections indicated with arrows); oscillator frequency as a function of timer state; regulatory logic of “temporal” model variant (timer state is a function of time); regulatory logic of “spatial” model variant (timer state replaced by a PI value, which is a function of distance). The frequency profile I have chosen scales the oscillator frequency by (1 *− e*^*−*2*τ*^)*/*(1 *− e*^*−*2^). **B,C:** Example simulation output (see Movie 1), for the parameter values and timepoints shown. Only one switch state is plotted. Note that faster oscillations (smaller *P*) produce shorter segments; a slower timer (larger *T*) produces a longer SAZ; faster elongation (larger *v*) produces longer segments and a longer SAZ. **D,E:** Simulation output compared between temporal variant (left, solid lines) and spatial variant (right, dotted lines), for a range of different parameter values. *L*_(SAZ)_ = number of cells where *τ >* 0, excluding signal-positive cells; *L*_(segment)_ = segment length; Δphase = oscillator phase difference between anterior and posterior of SAZ. The variants differ only in their response to elongation rate.

The temporal variant of the model involves a timer, an oscillator, and a switch. The timer represents cellular maturation from an undifferentiated (posterior SAZ) state to a differentiated (anterior SAZ) state. The oscillator represents a segmentation clock. Finally, the switch represents a binary choice between anterior and posterior segment polarity fates.

The timer is maintained in its initial state (1) by the posterior signal. Once this signal is removed, the timer decreases at rate 1*/T* until it reaches its ground state (0). When the timer reaches 0, it triggers a hand-over from the oscillator (expressed only when *τ >* 0) to the switch (expressed only when *τ* = 0), with segment-polarity fate (switch value) determined by the final oscillator level as it turns off (1 if *o >* 0.5; 2 otherwise). In addition, the frequency of the oscillator is modulated by the timer: a lower timer state causes slower oscillations^∗^. The regulatory interactions between the modules reflect, in simplified form, those between PSM TFs (the timer), the *hes*/*Notch* segmentation clock (the oscillator) and *mesp* segment polarity genes (the switch) in vertebrate somitogenesis, as inferred from genetic perturbations and functional analysis of enhancers [103–105, 202, 203]. (In arthropods, the interactions between SAZ TFs, oscillating genes, and segment-polarity genes are likely to be similar but slightly more complicated, as described in the section below.)

When simulated, the model generates typical “clock and wavefront” behaviour (Figure 3B,C, Movie 1). Because the state of the timer in a given cell is a function of the time since that cell last saw the posterior signal, elongation generates a dynamic gradient of timer state along the AP axis (*i*.*e*., a temporal coordinate system), which moves posteriorly in concert with the posterior signal. Fast, synchronous oscillations are maintained in the posterior of the embryo (where *τ* = 1); these narrow into anteriorly progressing kinematic waves where 0 *< τ <* 1. Finally, segment polarity expression is activated stably in a smooth A-P progression, as the “determination front” (*τ* = 0) moves posteriorly at a constant rate. As expected [39], the segmentation rate matches the oscillation period in the posterior of the embryo, and segment length depends jointly on the oscillator period and the elongation rate (*L*_seg_ = *P∗ v*). The inclusion of the timer in the model means that cells are destined to autonomously follow a particular temporal trajectory once they are relieved from the posterior signal. Thus, the spatial organisation of the SAZ is a reliable by-product of elongation, but plays no explicit role in cell regulation. Alternatively, segmentation dynamics could be under the control of a PI system: the SAZ would be associated with a typical lengthscale *λ*, and a cell’s behaviour would depend explicitly on its position within the SAZ field. This situation is represented by the “spatial” variant of the model, in which the timer is replaced by a state variable that is a function of a cell’s distance from the posterior signal (Figure 3A, right), and the same downstream effects are retained.

Although these model variants are conceptually distinct, they generally predict very similar behaviour (Figure 3D). In both cases, the progression of the wave-front is powered by elongation – either because elongation causes increasingly posterior cells to set their timers in motion at increasingly advanced timepoints (the temporal variant), or because elongation causes the whole SAZ field to retract along with the posterior signal (the spatial variant). Whether cell maturation is governed by a characteristic timescale (*T*, temporal variant) or by a characteristic lengthscale (*λ*, spatial variant), segment length is determined only by *P* and *v*. In addition, SAZ length scales linearly with either *T* or *λ*, as does the oscillator phase difference across the SAZ, which determines the number of SAZ stripes. The phase difference also scales with oscillator frequency in both models.

However, the two variants are affected differently by the elongation rate (Figure 3E, Movie 2). With a timer, SAZ length is proportional to elongation rate (*L*_SAZ_ = *T* ∗*v*), but with a PI system SAZ length depends (by definition) only on *λ*. Furthermore, with a timer the SAZ phase difference is essentially independent of the elongation rate, but with a PI system slower elongation is associated with an increasing number of stripes. A timer-based system therefore naturally predicts scaling of SAZ length with elongation rate and scaling of wave dynamics with SAZ length, while a PI-based system does not.

## 5 A clock and two timers

The next model (Figure 4) examines how a more complex segment-polarity pattern could be generated. Arthropods generate at least three states for every segmentation clock cycle (Figure 4C, middle), instead of just two states as in vertebrates (Figure 4C, top). Some groups of arthropods (insects, centipedes), pattern two segments for every oscillator cycle [14, 204], and must generate a pair of triplet repeats (*i*.*e*., six states; Figure 4C, bottom).^†^

**Figure 4:**
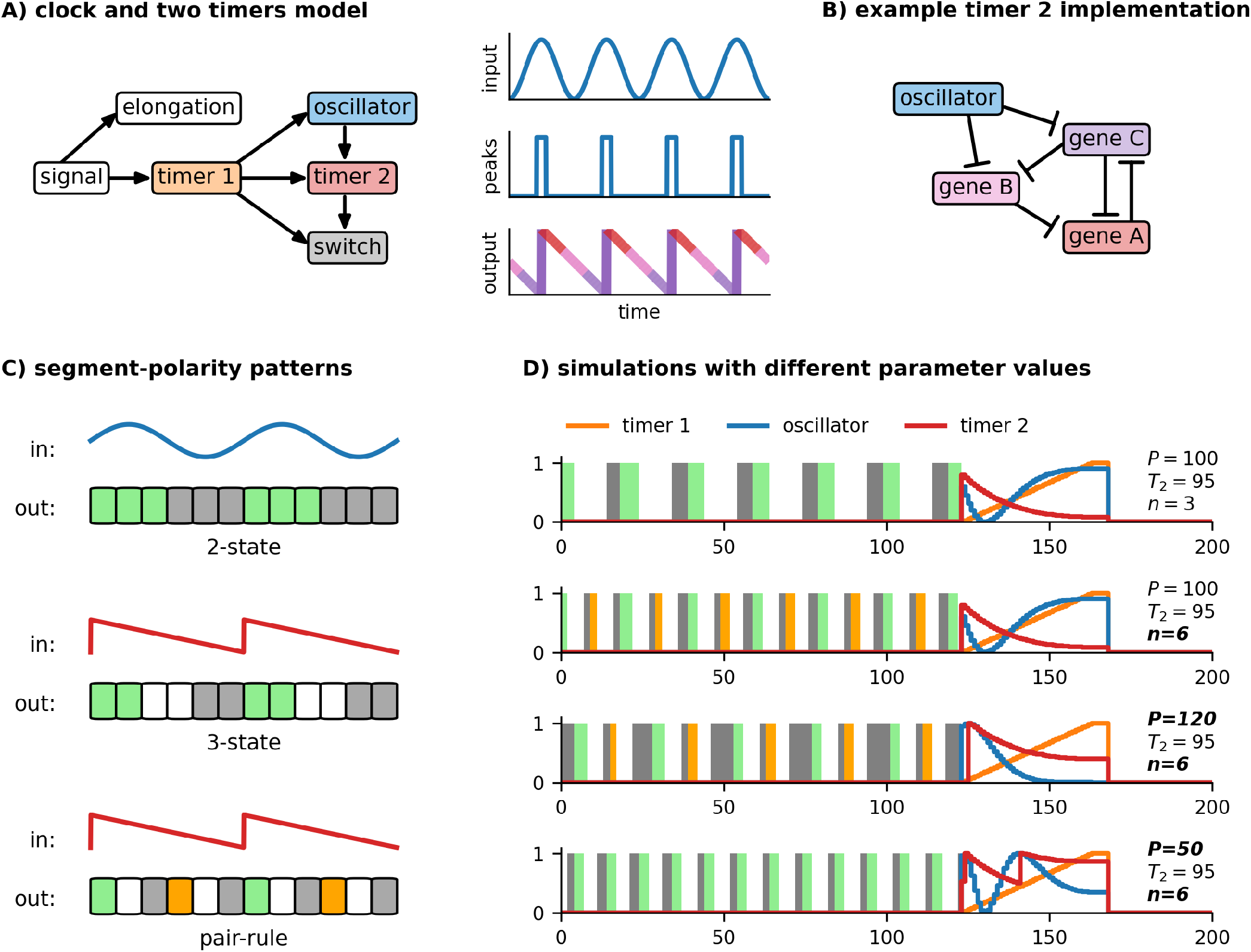
Clock and two timers model. **A:** left, diagram of model structure (dynamical modules shown as coloured boxes, other model features shown as white boxes, regulatory connections indicated with arrows). Right, cartoons of raw oscillator signal (top), oscillator signal digitised by a threshold (middle), sawtooth response of timer 2 to digitised signal (bottom). **B:** example network topology for timer 2. Genes A-C form an AC-DC circuit capable of driving an A→B→C sequence of expression [197]. High oscillator expression represses the two later-expressed genes (B and C), derepressing gene A and resetting the sequence to its initial state. **C:** Diagrams of different segment-polarity patterns. “3-state” and “pair-rule” patterns require polarised regulatory input. Green = anterior fate; grey = posterior fate; white = middle fate; orange = alternative anterior fate. **D:** Example simulation output (see Movie 3), for parameter values shown. *T*_2_ duration is the same in each case. The top two rows have the same oscillator period but different timer 2: switch mappings (3-state *vs* pair-rule). The bottom three rows all have the same timer 2: switch mapping (pair-rule) but different oscillator periods.

If cells are able to calculate “where” they are within an oscillation cycle (*i*.*e*., oscillation phase), it is theoretically possible to generate a segment-polarity pattern of arbitrary complexity, simply by changing the mapping between the input phase values and the output switch states. For example, two input thresholds are needed for a three-state pattern, and five thresholds are needed for a pair-rule pattern. The trivial way to implement this is to allow cells direct access to the oscillator phase variable that is “hidden” in the clock and timer model just described. Unambiguous phase determination would be possible, for example, if the segmentation clock was a multi-gene “ring oscillator” [205], because each portion of the phase would correspond to a unique combination of transcription factor concentrations. Under this scenario, the segment-polarity pattern would always scale with segment length.

However, suppose the core oscillator is based on direct *her* /*hes* gene autorepression, as is the case in vertebrates [206], and potentially arthropods as well [1]. Now there is no one-to-one mapping between oscillator output (*i*.*e*., Her/Hes protein concentration) and oscillator phase, because oscillator output both increases and decreases over the course of a cycle. (This is the motivation for hiding the phase variable from the other modules – in this scenario oscillator phase is implicit in the combination of oscillator transcripts and proteins present in a cell, but not determinable from the protein level alone.) For cells to unambiguously determine their position within a oscillator repeat, they must instead compute phase information from the oscillator’s *dynamics*.

Suppose an additional timer module is inserted into the clock and timer model, to mediate the interaction between the oscillator and the switch (Figure 4A, left). Let the original timer be “timer 1” (*τ*_1_), and the new timer “timer 2” (*τ*_2_). Like the oscillator, timer 2 can only be expressed where *τ*_1_ *>* 0, and so is similarly restricted to the SAZ. It also receives periodic input from the oscillator: when oscillator levels are at their peak (above a threshold *θ* = .995), timer 2 resets to its initial state (1), else it decreases at rate 1*/T*_2_ until it reaches its ground state (0). Timer 2 therefore measures the time since the oscillator was last at its peak, and produces a polarised sawtooth pattern downstream of symmetrical oscillator input (Figure 4A, right; note that the analog oscillator signal is effectively digitized to peak (ON) *vs* non-peak (OFF) as regards its downstream effects).

The inclusion of timer 2 in this model was inspired by the arthropod pair-rule gene network. In *Drosophila*, this network does indeed function downstream of a potential oscillator component (the *her* /*hes* gene *hairy* regulates, but is not significantly regulated by, other pair-rule genes), and upstream of the segmental pattern (pair-rule genes other than *hairy* directly regulate segment-polarity genes) [123, 207]. Several pair-rule genes oscillate in the arthropod SAZ [1, 92, 205, 208], and the documented interactions between them are consistent with the network topology shown in Figure 4B, which is one implementation of an oscillator-resettable timer [1, 123].^‡^

Combining an oscillator with a downstream timer allows for more flexible segment patterning than does the basic clock and timer model (Figure 4D, Movie 3). Because the switch is patterned by timer 2 rather than the oscillator, polarised (and potentially complex) segment-polarity patterns become easy to produce. In addition, because the oscillator and timer 2 each have their own characteristic timescale, the segment pattern is no longer forced to scale with segment length. If *T*_2_ is of similar magnitude to *P*, the state of timer 2 makes a good proxy for oscillator phase, and the resulting segment-polarity pattern will resemble the timer 2: switch mapping. However, if *T*_2_ *< P*, timer 2 will have to wait some time at 0 before the oscillator hits its next peak, and the final state in the timer 2: switch mapping will take up a larger proportion of the pattern than if the timescales were balanced. (If the oscillator cycles with double-segment periodicity, this effect will cause alternate segments to be different lengths, as seen in some scolopendrid centipedes.) Conversely, if *T*_2_ *> P*, timer 2 will always be reset by the oscillator before it reaches 0, and posterior segment-polarity states within the timer 2: switch mapping may not be produced. (In an extreme case, this effect could even change the periodicity of patterning from double-segmental to single-segmental.) Finally, if the oscillator and timer 2 happen to have different frequency profiles, the segment pattern will no longer be independent of the timescale of timer 1 (Supplementary Figure 2). This is because the phase relationship between the oscillator and timer 2 will vary across the length of the SAZ, and the longer it takes for a cell to exit the SAZ, the greater this divergence will become.

## 6 A clock and timer with feedback

Decoding the phase of symmetrical oscillations is thus one mechanism to generate a polarised spatial pattern. An alternative mechanism is to use the TST process to modify the oscillations themselves; periodicity is generated by symmetrical temporal oscillations as usual, but the spatial pattern recorded by the embryo is no longer a faithful copy of this input.

The motivation here is to create a sawtooth spatial output similar to that seen for CDH2 in nascent somites. CDH2 levels affect cell adhesion, and boundary formation is favoured where low and high levels of CDH2 directly abut [209]. Regulatory inputs from the segmentation clock establish a smooth gradient of CDH2 across each nascent somite [134], with sharp high-to-low transitions prefiguring the somitic fissures [133, 136]. Strikingly, an ectopic fissure forms in the middle of each somite in mouse *CDH2* mutants [210] (*i*.*e*., boundaries form at P-A *and* A-P transitions) suggesting that this sawtooth pattern is instructive for polarised morphogenesis.

A symmetrical temporal pattern can only be transformed into a polarised spatial pattern if the translation velocity varies over time. To illustrate the concept, suppose a wavefront is transcribing synchronous sinusoidal oscillations by freezing them in space (Figure 5). If the wavefront velocity is constant (Figure 5A), the spatial output pattern is just the temporal input pattern scaled by the velocity. In other words, because the wavefront moves at a uniform rate, each section of the temporal pattern is given equal weight in the spatial output.

**Figure 5:**
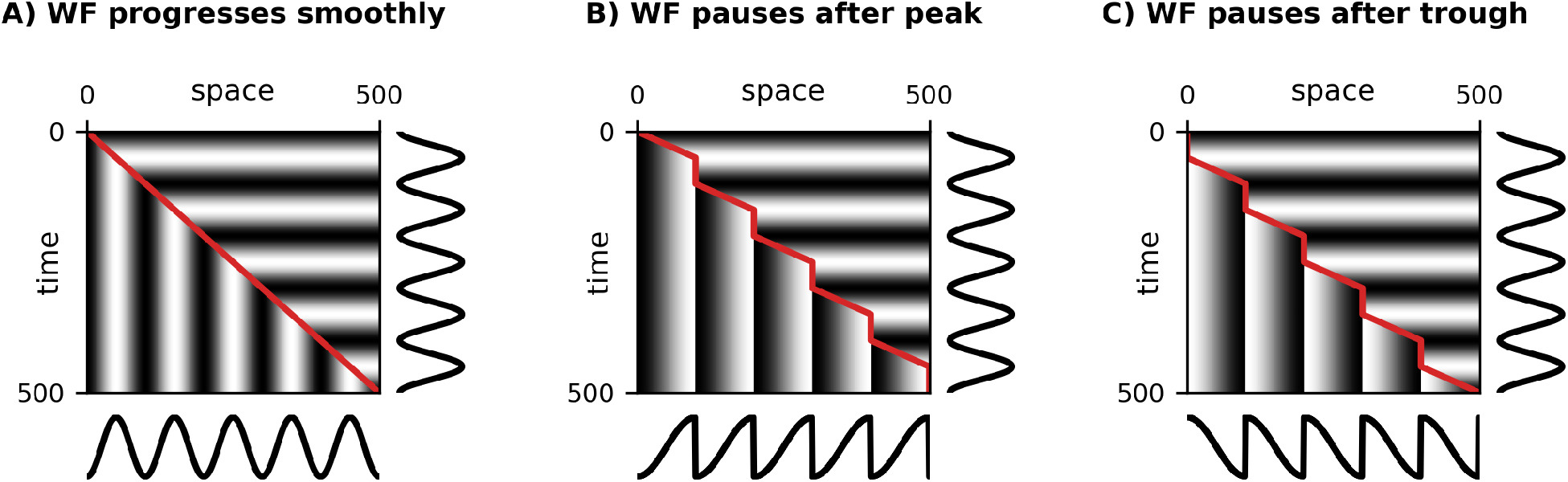
Effect of wavefront dynamics on time-space translation. **A:** kymograph for a simple clock and wavefront system, with a wavefront (red line) that moves at a constant velocity. The system faithfully reproduces the clock’s temporal signal in space. **B:** the wavefront progresses (at 2x speed) only while the clock signal is increasing from trough to peak, and pauses while it returns to the trough. A sawtooth pattern is recorded. **C:** the wavefront progresses (at 2x speed) only while the clock signal is decreasing from peak to trough, and pauses while it returns to the peak. A reverse sawtooth pattern is recorded.

If the wavefront velocity is variable, however, the output pattern will qualitatively change. If the wave-front moves faster or slower than average during some part of the oscillation cycle, the corresponding part of the input pattern will be stretched or compressed, respectively, in the output. If the wavefront pauses during part of the cycle, the corresponding section of the pattern will be skipped.

To extract a sawtooth pattern from the temporal signal, the wavefront would need to “transcribe” only half of the input pattern, stretch it to fill a whole segment length, and discard the rest – *i*.*e*., move at 2x speed for the first half of each oscillation cycle and pause for the second half (Figure 5B), or *vice versa* for reversed polarity (Figure 5C). (Note that the repeat length of the pattern is the same in all cases, but the lengthscale of the waves is doubled in Figure 5B,C as compared with 5A.) This patterning outcome requires that the wave-front velocity is strictly coordinated with both the oscillator period and the oscillator phase, suggesting that it could only occur in an embryo if there was feedback from the oscillator on the wavefront.

Sigificantly, pulsatile movement of the wavefront has been documented in vertebrates, and shown to be regulated by the segmentation clock [211–215]. In addition, zebrafish *her1* gene expression has been described as showing a sawtooth pattern and a double-segment lengthscale as it arrests, rather than the expected sinusoidal wave [201], suggesting that the non-uniform wavefront dynamics do indeed result in oscillation polarisation.

I now modify the clock and timer model to show how these effects could arise. Consider a stripped-down version of the clock and timer model that lacks the switch module (Figure 6A, left) and simply “freezes” the oscillator phase when the timer reaches 0, giving a clear readout of the transcribed pattern (Figure 6D). With this basic set-up, the wavefront moves at a constant velocity, the spatial output is isomorphic to the temporal input, and (as found earlier), neither the timer duration nor the oscillator frequency profile have any practical effect on the spatial output.

**Figure 6:**
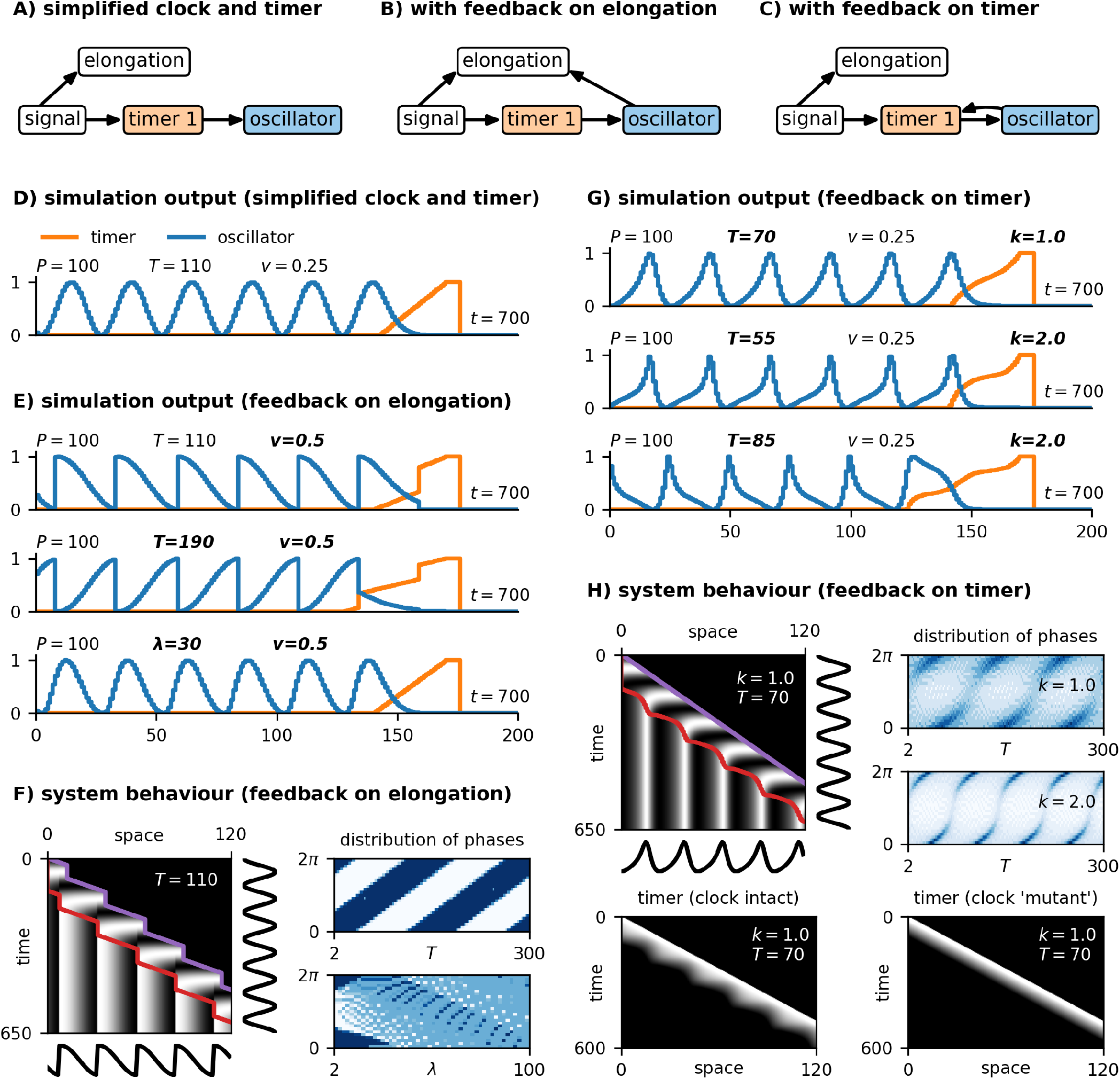
Clock and timer model with feedback. **A,B,C:** diagrams of simplified clock and timer model with and without feedback (dynamical modules shown as coloured boxes, other model features shown as white boxes, regulatory connections indicated with arrows). **D,E,G:** example simulation output (see Movies 4 and 5), for the model types and parameter values shown. **F:** left, kymograph of the simulation from the top row of E. Note that the posterior tip of the embryo (purple line) and the determination front (anterior extent of the SAZ, red line) both move in a stop-start manner. Right, plots showing the frequency distribution of recorded clock phases for a range of values of *T* (top, temporal variant) or *λ* (bottom, spatial variant); darker blue = higher frequency of phase within final output. (*I*.*e*., dark blue = over-representation, light blue = under-representation. For the model in A, the plot would show a uniform field of medium blue, because all oscillation phases are represented equally in the output.) **H:** top left, kymograph of the simulation from the top row of G. Note that the posterior tip of the embryo (purple line) moves smoothly, but the determination front (red line) moves in a pulsatile manner. Bottom left, kymograph showing timer 1 state during the same simulation. Bottom right, timer state from a simulation where the oscillator state is maintained at 0; note that the wavefront pulses are gone. Top right, plots showing the frequency distribution of recorded clock phases for a range of values of *T*, using a weak (top) or a strong (bottom) level of feedback on timer 1.

To affect the dynamics of the wavefront, the oscillator must modulate either the elongation rate or the timer. Suppose first that the oscillator affects elongation (Figure 6B): when the oscillator state in the posterior SAZ is below a threshold level (0.5) elongation proceeds as normal, but above this threshold elongation is suppressed. Axis extension now becomes coordinated with the segmentation clock, as does the movement of the wavefront, and only half of the temporal input pattern is transcribed (Figure 6E,F, Movie 4). The pattern that is finally recorded depends strongly on *T*_1_ (the timescale of timer 1) because this value in combination with the frequency profile determines which phases of oscillator expression will intersect with the stops and starts of the wavefront in the anterior SAZ.

If timer 1 is replaced by a PI field (recall Figure 3A, right) to produce a spatial variant of this model, embryo elongation and wavefront velocity show the same periodic dynamics as in the temporal variant. Interestingly, however, the PI field and frequency profile combine to produce a strong buffering effect on the SAZ stripes, so that the spatial output pattern only diverges strongly from temporal input pattern when the PI field is very short.

Alternatively, the feedback could act on the timer. Suppose that elongation rate remains constant but the oscillator affects the rate at which the timer state decreases (Figure 6C). For example, let the timer rate be proportional to *e*^*−ko*^, where *o* is oscillator level and *k* is a feedback strength parameter; the timer will decrease more slowly in a given SAZ cell when its oscillator level is high. The wavefront velocity now becomes decoupled from the elongation rate, and the spatial output pattern is again affected (Figure 6G,H, Movie 5). As just described for the model with feedback on elongation, the spatial pattern that is eventually recorded depends strongly on *T*_1_, as well as on the strength of the feedback.

## 7 A clock and three timers

I now turn back to the clock and two timers model, and modify it to allow for axial variation and finite body length. Suppose a third timer (“timer 3”), representing Hox and/or gap gene dynamics, is added downstream of timer 1 (Figure 7A). Within the SAZ, timer 3 decreases from its initial state (1) to its ground state (0) at rate 1*/T*_3_. Outside the SAZ, timer 3 remains stable. As a result of TST, the timer 3 value of a mature segment provides a proxy measure of its axial position and can be used to specify its identity. To modulate segment size and trigger segmentation termination, timer 3 additionally needs to affect the wavefront velocity, as discussed above. This implies that it should modulate the elongation rate, and potentially timer 1 as well.

**Figure 7:**
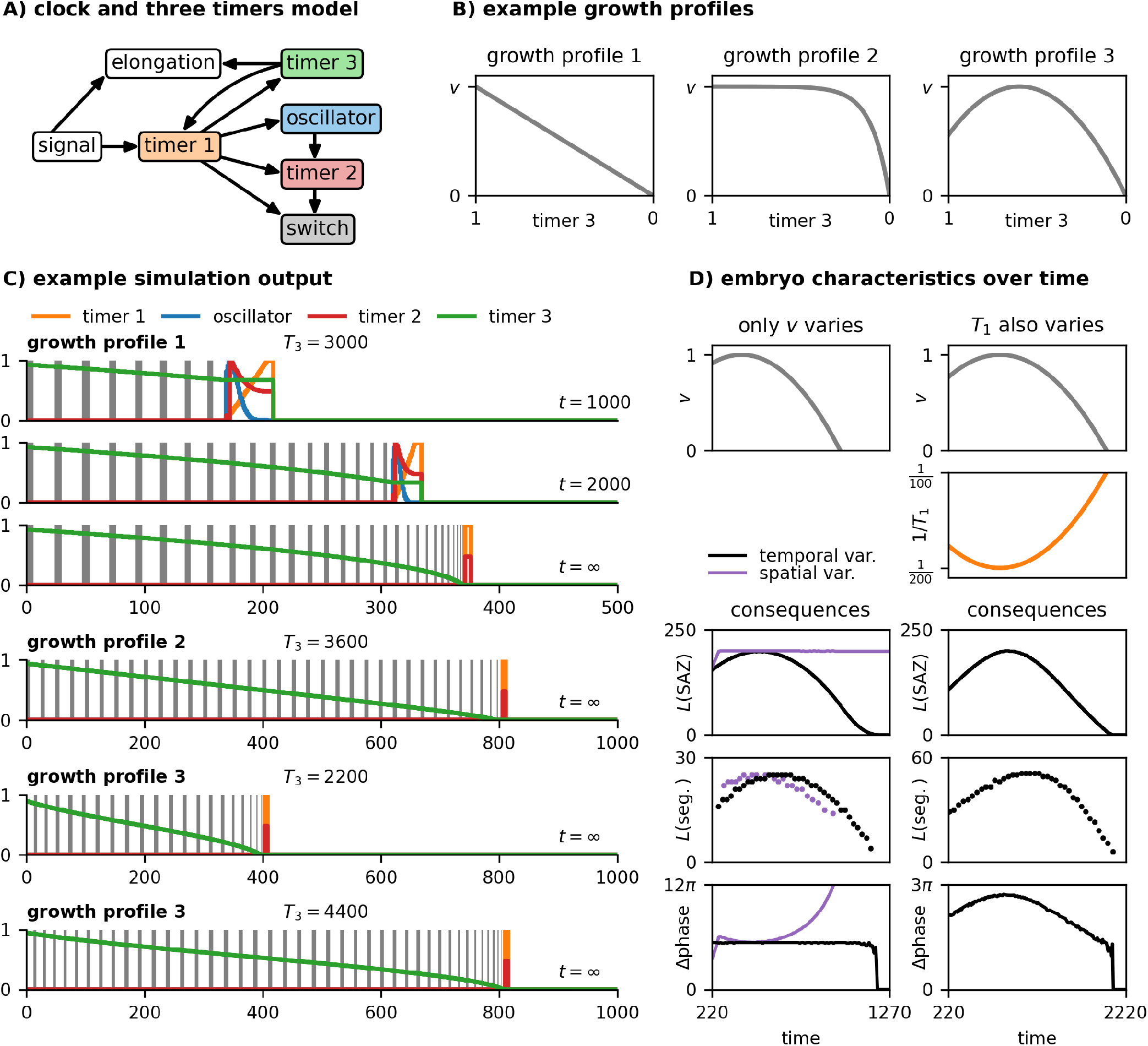
Clock and three timers model. **A:** diagram of model structure (dynamical modules shown as coloured boxes, other model features shown as white boxes, regulatory connections indicated with arrows). **B:** example growth profiles; note reversed x-axis. **C:** example simulation output (see Movie 6), for parameter values, growth profiles, and timepoints shown. Temporal model variant used. “*t*=*∞*” indicates that the final segment pattern has been reached. **D:** summary statistics over time for a scenario with a changing growth rate (left, see Movie 7) and a scenario with both a changing growth rate and a changing timer 1 decrease rate (right, see Movie 10). For the first scenario, simulation results from both the temporal model variant (black) and the spatial model variant (purple) are shown. The time axes start at 220 to crop out the early stages of the simulations, where length of the growing embryo is smaller than the length of a mature SAZ.

Suppose the elongation rate depends on the state of timer 3, with the mapping between *τ*_3_ and *v* defined by a particular “growth profile” (Figure 7B). The growth profile can be of arbitrary shape but always falls to 0 when *τ*_3_ = 0, thereby terminating axis extension and segment addition. Regardless of whether the temporal or spatial variant of this model is used (recall Figure 3A, right), the segment size distribution along the AP axis will mirror the shape of the growth profile, and the final length of the axis (and therefore the final segment number) depends on the magnitude of *T*_3_ (Figure 7C, Movie 6).

Other aspects of system behaviour differ between the two variants (Figure 7D left, Movie 7). As found earlier, SAZ length scales with elongation rate when the SAZ is patterned by a timer. Accordingly, with the temporal variant, the length of the SAZ over time mirrors the growth profile, albeit with an adaptation time lag that depends on the magnitude of *T*_1_. In addition, because the phase difference across the SAZ is independent of the elongation rate, the number of waves within the SAZ remains constant, the lengthscale of the pattern adapting automatically to the changing SAZ length. In contrast, the behaviour of the spatial variant is almost the opposite: the length of the SAZ is constant over time (because it is defined by *λ* rather than *T*_1_∗*v*), whereas the phase difference across the SAZ falls as the elongation rate rises and rises as the elongation rate falls (because the elongation rate determines how long each cell remains within the SAZ). The most striking difference between the temporal and spatial variants occurs after elongation stops: in the temporal variant, the SAZ shrinks and forms segments until it is completely exhausted; in the spatial variant, the SAZ persists indefinitely and accumulates a huge build up of increasingly crowded-together stripes^§^.

Clearly, the behaviour of the temporal variant is closer to the behaviour of real embryos than is the behaviour of the spatial variant. Indeed, with the temporal variant, the time lag required for a changing elongation rate to affect the length of the SAZ, and eventually the length of new segments, reproduces (and potentially explains) a scaling phenomenon recently discovered in zebrafish, whereby the length of new segments is proportional to past SAZ length but not to current SAZ length [47] (see Supplementary Figure 3).

The temporal variant does diverge from experimental observations, however, by generating an SAZ phase difference that is entirely constant over time. In real embryos, this parameter peaks at the beginning or middle of segmentation before undergoing a gradual decline [149]. More realistic dynamics can be recovered if timer 3 additionally modulates timer 1: for example, if SAZ cells are made to mature faster towards the end of segmentation, the SAZ phase difference becomes accordingly smaller (Figure 7D right, Movie 10). Other feedback effects can also be incorporated into the model. For example, were timer 3 to reduce the *initial state* of timer 1 (as opposed to its timescale), the oscillation period in the posterior of the embryo would decrease as dictated by the oscillator frequency profile, mimicking the slower oscillations that are observed towards the end of segmentation or when Wnt levels are reduced [87, 104, 150, 173].

## 8 Three timers and no clock

Finally, I show how the clock and three timers model can be modified to produce simultaneous rather than sequential patterning.

Kymographs from a typical clock and three timers simulation are shown in Figure 8D, top. Timer 1 (left panel) provides a temporal framework; timer 3 and the oscillator (middle panel) generate state differences between cells; and timer 2 and the switch (right panel) together transduce these differences into stable fates. The origins of the final pattern lie, therefore, in the dynamics of the oscillator and timer 3: the oscillator generates a periodic pattern that is transduced into different segment polarity fates, while timer 3 generates a monotonic pattern that is transduced into region-specific differences.

**Figure 8:**
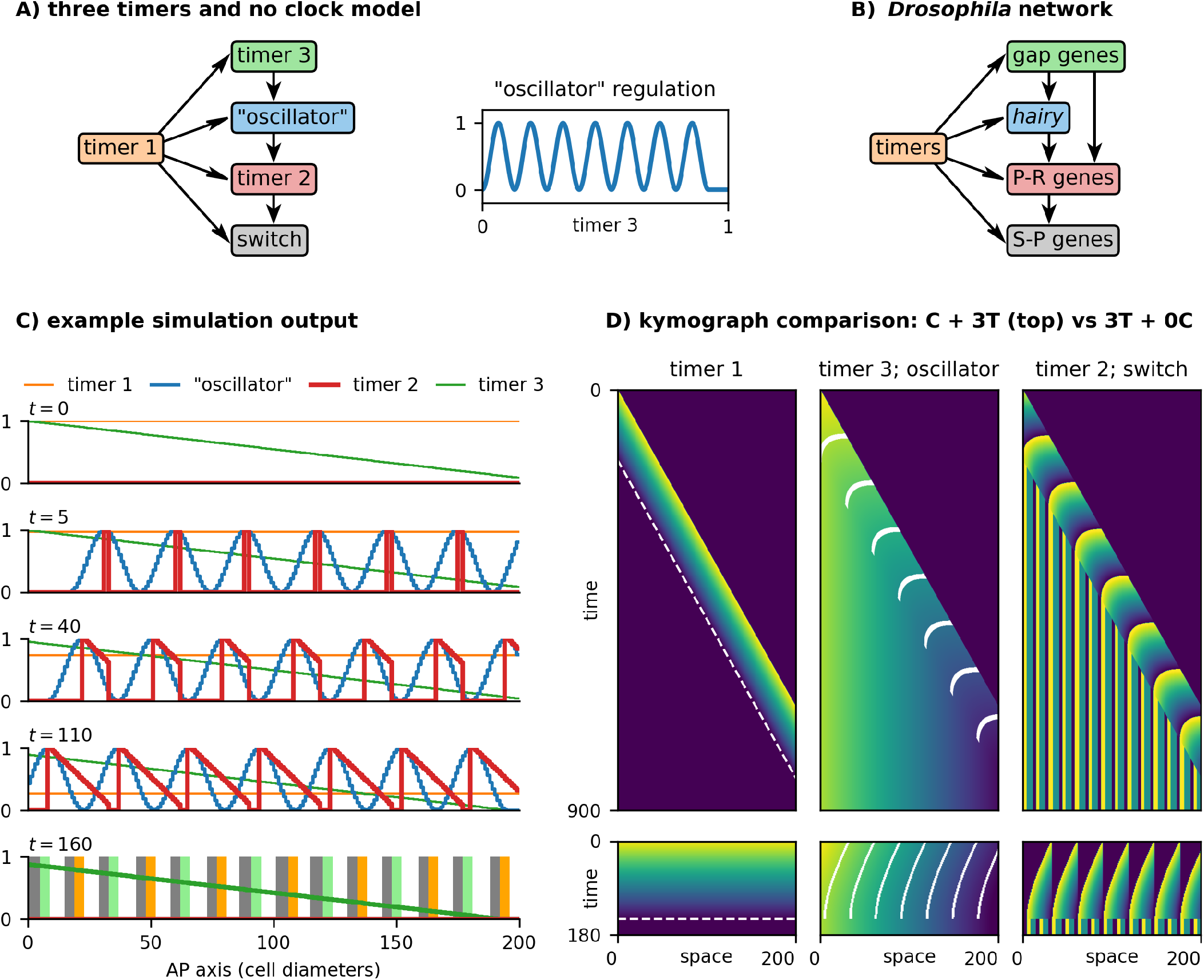
Three timers and no clock model. **A:** left, diagram of model structure (dynamical modules shown as coloured boxes, regulatory connections indicated with arrows). Right, complex mapping between timer 3 input state and “oscillator” output state. **B:** diagram showing the topology of the *Drosophila* segmentation network. “P-R” = pair-rule; “S-P” = segment-polarity. **C:** simulation output (see Movie 11) at five timepoints. Top row shows initial conditions; bottom row shows final, stable state. **D:** kymographs comparing “clock and three timers” output (top) with “three timers and no clock” output (bottom, same data as D). Dashed white lines overlaid on timer 1 output (left) indicate the determination front. Solid white lines overlaid on timer 3 output (middle) mark oscillator peaks. Timer 2 values (right, SAZ pulses) are converted into switch states (right, stable domains) at the determination front. For each module, yellow = 1 and purple = 0. Time axes are to scale. See also Movie 11.

In essence, time is used to generate sequences of *o* and *τ*_3_ states, which the rest of the system then translates into spatial patterns. The overall mechanism is elegant, but not very efficient – as the kymographs demonstrate, the actual time that each segment is actively undergoing patterning is short, with a much longer time spent waiting around for anterior or posterior segments to undergo maturation. The process would finish much faster if segment patterning could be parallelised.

Suppose an embryo is initialised at its final length, and, in the absence of a PSC, timer 1 counts down in all cells simultaneously. Without axial elongation, spatial differences in *o* and *τ*_3_ are no longer generated automatically, and must be introduced by other means. Since the required timer 3 pattern is monotonic and the required oscillator pattern is not, it seems reasonable to assume that the timer 3 pattern is the easier to create *de novo*. I therefore build a monotonic gradient of timer 3 states into the embryo’s initial conditions, analogous to the early expression of *Drosophila* gap genes that is established directly downstream of maternal gradients [157]. I then use this new source of spatial information to recover stripes of oscillator expression, by changing the regulation of the oscillator module: it no longer has autonomous dynamics, but is instead a function of timer 3 (Figure 8A, left; compare Figure 7A). The complex, periodic mapping between timer 3 input and “oscillator” output (Figure 8A, right) is analogous to the way that the *Drosophila* embryo generates a serial array of *hairy* stripes from gap gene inputs, thanks to the evolution of various stripe-specific enhancers [182, 216].

Simulating the new model (Figure 8C, Movie 11) produces the same final output as the clock and 3 timers model, in just a fraction of the time. As can be appreciated from the kymographs of the simulation (Figure 8D, bottom), although the tissue-level dynamics are different from the clock and three timers model, its intracellular dynamics are preserved. As timer 3 gradually ticks down in each cell (albeit only through a fraction of its full dynamic range), its monotonic spatial pattern marches anteriorly across the tissue, dragging the “oscillator” pattern with it to produce spatial waves. Because these waves recapitulate the dynamical output of an intact oscillator, the stripe peaks are still able to generate temporal gradients of timer 2 expression in their wakes, just as they would in the SAZ. Finally, shortly after the timer 2 pattern is completed, timer 1 triggers the “determination front” in every cell simultaneously, stabilising the entire pattern via the activation of the switch.

Comparing the three timers and no clock model (Figure 8A) to the *Drosophila* segmentation hierarchy (Figure 8B) [1, 217] reveals that their structure is almost identical, except that the gap genes also provide considerable direct input into non-*hairy* pair-rule genes. In *Drosophila*, the pair-rule genes (timer 2) are patterned partially by stripe-specific enhancers (PI) [182] and partially by cross-regulatory dynamics (TST) [122, 123, 193], rather than purely by TST as in the model. The model could indeed be sped up even further if timer 3 additionally helped to establish timer 2, because most of the time required to establish the final pattern is taken up by timer 2 dynamics. Indeed, it is notable that gap gene-mediated patterning is most important for *Drosophila* pair-rule genes expressed “earlier” in the pattern repeat (assuming a repeat stretches from one *hairy* stripe to the next), while cross-regulation dominates for those expressed “later”.

## 9 Discussion

In this article, I have used a simple modelling framework to explore the tissue-level consequences of interlinked cell-autonomous dynamical processes in the context of segmentation. I have compared the ramifications of temporal *vs* spatial regulation of the SAZ, considered the various spatial outputs that can be generated downstream of an oscillation, and examined the evolutionary relationship between sequential and simultaneous segment patterning. Here, I discuss the specific implications of each model, the developmental significance of TST, and the advantages and disadvantages of the “dynamical module” approach.

### 9.1 Temporal or spatial patterning of the SAZ?

SAZ patterning has been thought about in a number of different ways. One conceptual division concerns whether the anterior and posterior ends of the SAZ are functionally linked. In some models (usually informal), the anterior boundary of the SAZ is determined by the rate of tissue maturation, the posterior boundary of the SAZ is determined by the rate of axial elongation, and the two independent processes must be coordinated (“balanced”) to maintain an appropriate SAZ length [111, 150]. In others, the rate of axial elongation also determines the rate of tissue maturation [39, 140].

The “temporal” and “spatial” model variants I have investigated here are both of the latter type. However, the role played by elongation rate differs between them. In the spatial variant, the size of the SAZ and its maintenance over time are both independent of elongation; if elongation stops, tissue maturation stops and the SAZ remains the same size. In the temporal variant, SAZ size scales with elongation rate; if elongation stops, tissue maturation continues and the SAZ shrinks until it eventually disappears.

This difference has significant implications for segmentation dynamics, particularly when the elongation rate and/or other system parameters vary over time. Temporal patterning provides a mechanistic explanation for the scaling of SAZ size with embryo size [47] and predicts constant [44] or smoothly-varying [149] wave patterns, while spatial patterning does neither. In addition, temporal patterning is very sensitive to variation in elongation rate whereas spatial patterning buffers it; these properties could be advantages or liabilities depending on whether such variation plays instructive or destructive roles in the system. Finally, temporal patterning significantly reduces the interpretational burden of the cell (because its state trajectory can be partially autonomous), and also removes the expectation that the determination front should necessarily correspond with a particular signalling threshold [48].

The strict opposition between temporal (TST) and spatial (PI) patterning presented here is artificial; in reality, an SAZ will be patterned by both. Experiments that characterise cell state trajectories under a range of constant conditions could help quantify their relative contributions. Similar experiments combined with misexpression of (for example) Hox and gap genes would be informative about the mechanistic origins of axial variation.

### 9.2 From oscillations to polarisation

The problem of segment polarisation and the inadequacy of a two-state prepattern was recognised by Meinhardt even before the expression of any segment-polarity gene had been visualised [218, 219]. Here, I have demonstrated three strategies by which a sinusoidal temporal input can be used to specify a polarised spatial pattern: (1) using a timer network to calculate oscillation phase; (2) having the oscillator regulate the rate of elongation; and (3) having the oscillator regulate the rate of cell maturation.

The first strategy is consistent with, and inspired by, the cross-regulatory interactions between the pair-rule genes in *Drosophila* [1, 123], which likely evolved in a sequential segmentation context [92, 123, 220]. However, as yet relatively little is known about arthropod segmentation clocks and the way they regulate their downstream targets [1]. Characterising these networks in sequentially segmenting model species such as *Tribolium* (beetle) [14, 205], *Oncopeltus* (hemipteran bug) and *Parasteatoda* (spider) is a research priority.

A more general point illustrated by the timer strategy is that the phase relationship of linked dynamical processes will change along the SAZ if they do not share a common frequency profile. This has been discussed in the context of the vertebrate segmentation clock, which combines direct *her* /*hes* autorepression and indirect auto repression via Notch signalling. Modelling suggests that the time delays in the two feedback loops should scale similarly along the SAZ, to avoid the oscillations being extinguished [200, 221]. For other cross-regulating processes, altered phase relationships along the SAZ have been clearly documented, and the phenomenon appears to be functionally important for segment patterning [211, 222, 223].

The other two strategies involve generating a pulsatile rather than smooth progression of the wave-front, so that only certain phases of the oscillations are recorded. Rhythmic SAZ morphogenesis has been described in insects [67], where *Toll* genes regulate convergent extension downstream of the clock [66]. In vertebrates, cell proliferation has been found to occur preferentially at the trough phase of an oscillation [148], providing another mechanism by which elongation and oscillation could be coordinated. *her* /*hes* oscillations have also been shown to delay cell differentiation in cultured cells [224], while wavefront progression appears to be modulated by the segmentation clock in zebrafish [212, 213]. Combined with the sawtooth arrest dynamics of zebrafish oscillations [201], these observations suggest that the oscillator frequency profile might play a functional role in patterning somite morphogenesis.

Finally, it is significant that the frequency profile plays an explicit role in all three of the polarisation mechanisms I have described. Progressively narrowing travelling waves (a consequence of the frequency profile) are one of the most visually striking features of segmenting systems, but in most segmentation models they lack an instructive function [16, 39, 47, 225].

### 9.3 From sequential to simultaneous

The final model (three timers and no clock) demonstrates the considerable dynamical and regulatory continuity between sequential and simultaneous modes of segmentation. The central idea is that regionalisation processes, ancestrally parallel to and independent of segment patterning, provided a rich source of spatial cues that could be exploited to compensate for the loss of autonomous oscillations and the SAZ framework [1, 123, 226]. Thus similar expression relationships are observed between gap genes and pair-rule genes in sequentially segmenting and simultaneously segmenting species, but stripe formation is only governed by gap inputs in the latter.

Importantly, the preservation of the regulatory machinery downstream of the “oscillator” means that the regulatory links between the regionalisation system (gap genes) and the segment patterning system (pair-rule genes) need not be extensive, so long as the ancestral gap gene expression dynamics are preserved. At minimum, the gap gene pattern need specify only the start location of each pattern repeat (solid white lines in Figure 8D, bottom), and cross-regulation will fill in the rest. In *Drosophila*, the regulatory links between the gap genes and pair-rule genes are considerably more far-reaching than this [182], and some of the presumed ancestral cross-regulation between the pair-rule genes has apparently been lost [123] Indeed, gene expression shifts are subtle in most of the embryo, and absent up to around the second or third pair-rule repeat [166, 189, 190]. However, expression shifts are larger and more extensive in other simultaneously segmenting species [183, 227–231], and these extant forms may resemble transitional forms within the *Drosophila* lineage. Comparative analysis of pair-rule gene regulation in these species will be instructive.

### 9.4 Time-space translation in developmental systems

The central idea of TST is that spatial information can be specified by the combination of a timer and a velocity, because distance = velocity ∗time. Many of the patterning mechanisms in this article depend on the intersection of a moving posterior signal (a velocity) with autonomous cellular processes (time). Most of the patterning burden falls on the latter, whose regulatory logic, rates, and time delays would be implicitly encoded in the sequences of enhancers, exons, and UTRs. Thus, the interaction between inherited information and a directional morphogenetic process enables a complex pattern to emerge from a simple initial state.

In the somitogenesis literature, it has sometimes been stated that the segmentation clock determines *when* patterning occurs, while the wavefront determines *where* patterning occurs [40, 111]. In reality, given that the frequency profile generates spatial waves from the oscillations, and that these may in turn feed back on the wavefront, the division of labour is not this distinct. It is more useful to think about spatial information being split between various different processes, and repeatedly redistributed between them.

The final model I present provides a particularly clear example. For one thing, the supposed temporal *vs* spatial roles of “clock” (oscillator) and “wavefront” (timer 1) have been reversed. More importantly, the spatial information in the system is twice converted between different forms. First, the initial spatial pattern of timer 3 combines with the temporal dynamics of timer 3 to produce posterior-to-anterior waves (distance/time = velocity). This results in waves of “oscillator” expression, which then combine with the temporal dynamics of timer 2 to produce the spatial inputs to the switch (velocity∗ time = distance).

Thus, while there is an effective mapping between the timer 3 pattern at the beginning and the segment-polarity pattern at the end, the specific nature of the mapping depends on various dynamical rates. In *Drosophila*, this means that accurate and precise prediction of pair-rule gene outputs from gap gene inputs [232] is to be expected, but it need not imply that pair-rule gene regulation involves corresponding detail in signal decoding. As described in the previous section, the dynamics of patterning mean that only a subset of pair-rule stripes need to be positioned exactly, and selection on the gap gene system is likely to have adapted it specifically to this task [1].

### 9.5 Combining patterning strategies

In most of the models I have presented, the only intercellular signalling process is the Boolean posterior signal. The cells follow autonomous dynamical trajectories and produce coherent spatial patterns only because of their spatial ordering and temporal precision. This framework highlights the patterning capabilities of TST but is obviously incongruous with noisy, three-dimensional biology. In reality, the mechanisms presented here would be integrated with local cell communication, longer-range signalling, and morphogenetic feedback.

For example, Notch signalling-mediated self-organisation is required to counteract the oscillator desynchronisation produced by gene expression noise, division-induced expression delays, and cell rearrangement [37, 69, 233, 234]. Beyond simply preventing oscillations from deteriorating, the process also influences their period, amplitude, and frequency profile [33, 39, 118, 144, 235–239]. It will be necessary to study the interactions between self-organisation, PI, and TST mechanisms to properly understand the logic and reproducibility of developing systems.

Even the *Drosophila* blastoderm is a case in point. The establishment of the pattern of gap gene expression, which is impressively robust to variation in egg size and gene dosage [195, 240], seems to involve both PI and self-organisation mechanisms. The Bicoid concentration gradient is clearly instructive [241], but there is not a strict correspondence between Bicoid concentration and output gene expression to support strongly interpretational pattern formation [194, 195]. Indeed, diffusion of gene products between nuclei and cross-regulation between early gradients point to important reaction-diffusion effects [242–246]. Adding in the TST involved in segment patterning, it seems that all three patterning mechanisms are cooperating before complex morphogenesis has even begun.

### 9.6 The nature and evolution of dynamical modules

The “dynamical modules” of the models represent the expression dynamics of gene regulatory networks, potentially incorporating other autonomous processes such as signal decay. The focus of the models is on how different sets of dynamical processes might interact with or modulate each other, and how this could generate or transform spatiotemporal patterns.

There is an undeniable need for experimental and modelling studies that link the dynamical behaviours of networks to their topology and quantitative features [18, 197, 231, 247–251], something my study does not attempt to do. However, there are also advantages to using such a high level of abstraction. First, it focuses attention on the functional role of a dynamical behaviour, rather than its specific molecular implementation, making it easier to recognise general results. Second, a potentially complex process can be represented by a single parameter value (such as an effective timescale), rather than by a large number of regulatory interactions and rates, some of which may be experimentally inaccessible.

Beyond simple convenience, the dynamics of a process may be more conserved and/or functionally relevant than its mechanistic basis. Evolutionary systems drift [252] and local adaptation may cause the regulatory logic of homologous processes to diverge, even while their output is maintained by selection. See, for example, the differences in *her* /*hes* gene regulation across different vertebrate segmentation clocks, even though the genes’ oscillatory dynamics and down-stream functions are essentially unchanged [174, 206, 253–255]. At the same time, unrelated processes may share similar functions owing to convergent dynamics [256, 257] — different clock and wavefront systems (including arthropod segmentation and vertebrate somitogenesis) are a case in point [258, 259].

Finally, certain “modules” may be strikingly conserved across evolution, and perform intriguingly pleiotropic developmental roles. The same temporal sequence of gap gene expression is seen not only in simultaneous and sequential segmentation as discussed here, but also in the patterning of neuroblasts [260–263]. Hes oscillations, too, are seen in the nervous system [264– 266] and various other cellular contexts [224, 267, 268], while Hox gene dynamics are conserved across bilaterian AP axes [153, 162, 269] and reiterated in the vertebrate limb [270, 271]. Studying the origin, mechanistic basis, and evolutionary cooption/adaptation of dynamical modules is likely to provide insight into both micro- and macro-evolutionary change.

## Supplementary Materials

- **Supplementary Document**: model and simulation details.
- **Supplementary Figure 1**: summary diagrams of all models.
- **Supplementary Figure 2**: additional results from the clock and two timers model.
- **Supplementary Figure 3**: additional results from the clock and three timers model.
- **Supplementary Movie 1**: simulations of the clock and timer model, corresponding to Figure 3B,C.
- **Supplementary Movie 2**: simulations comparing temporal and spatial variants of clock and timer model.
- **Supplementary Movie 3**: simulations of the clock and two timers model, corresponding to Figure 4D.
- **Supplementary Movie 4**: simulations of the clock and timer model with feedback (on elongation), corresponding to Figure 6D,E.
- **Supplementary Movie 5**: simulations of the clock and timer model with feedback (on the timer), corresponding to Figure 6D,G.
- **Supplementary Movie 6**: simulations of the clock and three timers model, corresponding to Figure 7C.
- **Supplementary Movie 7**: simulations comparing temporal and spatial variants of the clock and three timers model, corresponding to Figure 7D, left.
- **Supplementary Movie 8**: simulations corresponding to Supplementary Figure 3B.
- **Supplementary Movie 9**: simulations corresponding to Supplementary Figure 3C.
- **Supplementary Movie 10**: simulation of the clock and three timers model, corresponding to Figure 7D, right.
- **Supplementary Movie 11**: simulations comparing the three timers and no clock model with the clock and three timers model, corresponding to Figure 8C,D.

## Acknowledgments

I thank Michael Akam, Matthew Benton, Rosa Martinez-Corral and Timothy Harden for comments on the manuscript.

## Footnotes

§ ^∗^The mechanistic basis for the frequency profile is still unclear, but seems to involve cell state dependent time delays [200]. Note that empirically characterised frequency profiles do not decrease smoothly to 0 as shown in Figure 3A [20, 115, 201], but the discrepancy is not important for my conclusions.

† This is known as “pair-rule” patterning, or double-segment periodicity. *N*.*B*., the “pair-rule gene” class of transcription factors are not universally expressed in pair-rule patterns, despite the name.

‡ More work is needed to characterise pair-rule gene cross-regulation in sequentially segmenting arthropods, and to understand whether pair-rule gene oscillations are autonomously generated, entrained by an upstream oscillator, or both. By characterising extant oscillation networks, the history of the arthropod segmentation clock may become clearer.

§ The model variants also show distinct responses to abrupt changes in elongation rate (as might be produced by experimental perturbations); see Supplementary Figure 3 and Movies 8 and 9.

