## Supplementary Document for "Time and Space in Segmentation"

### Model Details

| Symbol | Meaning |
|---|---|
| $t$ | time |
| $x$ | cell position |
| $W$ | signal |
| $\tau_{(1)}$ | timer (1) |
| $\phi$ | oscillator phase |
| $o$ | oscillator level |
| $s$ | switch |
| $T_{(1)}$ | duration of timer (1) |
| $\lambda$ | SAZ lengthscale |
| $P$ | oscillator period |
| $\theta$ | oscillator threshold |
| $k$ | feedback strength parameter |
| $L$ | embryo length |
| $v$ | elongation rate |
| $r$ | signal range |
| $f(\tau_2)$ | regulatory function for $s$ |
| $g(\tau_3)$ | growth profile |
| $h(\tau_3)$ | $T_1$ (or $\lambda$) scaling function |
| $a(\tau_3)$ | regulation function for $o$ |
| $b(x)$ | function that gives initial state of $\tau_3$ |

Table 1: Table of model symbols.

#### 1 Clock and Timer

Initial conditions: $\tau_{(0)} = 1$, $\phi_{(0)} = 0$, $o_{(0)} = 0$, $s_{(0)} = 0$.

$$\tau_{(t+1)} = \max\left\{\tau_{(t)} - (1 - W)/T,\ 0\right\} \tag{1.1}$$

$$\phi_{(t+1)} = \begin{cases} o_t + \left(\frac{1-e^{-2\tau_{(t)}}}{1-e^{-2}}\right) 1/P \mod 1, & \text{if: } \tau_{1(t)} > 0 \\ 0, & \text{otherwise.} \end{cases} \tag{1.2}$$

$$o_{(t+1)} = \frac{\sin(2\pi(\phi_{(t+1)} - \frac{1}{4})) + 1}{2} \tag{1.3}$$

$$s_{(t+1)} = \begin{cases} 2, & \text{if: } (s_{(t)} = 2) \cup ((\tau_{(t)} = 0) \cap (o_{(t)} \geq .5)) \\ 1, & \text{if: } (s_{(t)} \neq 2) \cap (\tau_{(t)} = 0) \cap (o_{(t)} < .5) \\ 0, & \text{otherwise.} \end{cases} \tag{1.4}$$

#### 2 Clock and two Timers

Initial conditions: $\tau_{1(0)} = 1$, $\phi_{(0)} = 0$, $o_{(0)} = 0$, $\tau_{2(0)} = 0$, $s_{(0)} = 0$.

$$\tau_{1(t+1)} = \max\{\tau_1(t) - (1-W)/T_1,\ 0\} \tag{2.1}$$

$$\phi_{(t+1)} = \begin{cases} o_t + \left(\frac{1-e^{-2\tau_{(t)}}}{1-e^{-2}}\right) 1/P \mod 1, & \text{if: } \tau_{1(t)} > 0 \\ 0, & \text{otherwise.} \end{cases} \tag{2.2}$$

$$o_{(t+1)} = \frac{\sin(2\pi(\phi_{(t+1)} - \frac{1}{4})) + 1}{2} \tag{2.3}$$

$$\tau_{2(t+1)} = \begin{cases} 1, & \text{if: } (\tau_{1(t)} > 0) \cap (o_{(t)} > \theta) \\ \max\left\{\tau_{2(t)} - \left(\frac{1-e^{-2\tau_{1(t)}}}{1-e^{-2}}\right) 1/T_2,\ 0\right\}, & \text{if: } (\tau_{1(t)} > 0) \cap (o_{(t)} \leq \theta) \\ 0, & \text{otherwise.} \end{cases} \tag{2.4}$$

$$s_{(t+1)} = \begin{cases} s_{(t)}, & \text{if: } s_{(t)} > 0 \\ f(t_{2(t)}), & \text{if: } (s_{(t)} = 0) \cap (\tau_{1(t)} = 0) \\ 0, & \text{otherwise.} \end{cases} \tag{2.5}$$

#### 3 Clock and Timer with feedback

Initial conditions: $\tau_{(0)} = 1$, $\phi_{(0)} = 0$, $o_{(0)} = 0$.
Use (3.1a) and (3.4a) for feedback on the elongation rate, and (3.1b) and (3.4b) for feedback on timer 1.

$$\tau_{(t+1)} = \max\left\{\tau_{(t)} - (1-W)/T,\ 0\right\} \tag{3.1a}$$

$$\tau_{(t+1)} = \max\left\{\tau_{(t)} - e^{-ko_{(t)}}(1-W)/T,\ 0\right\} \tag{3.1b}$$

$$\phi_{(t+1)} = o_t + \left(\frac{1-e^{-2\tau_{(t)}}}{1-e^{-2}}\right) 1/P \mod 1 \tag{3.2}$$

$$o_{(t+1)} = \frac{\sin(2\pi(\phi_{(t+1)} - \frac{1}{4})) + 1}{2} \tag{3.3}$$

$$v_{(t+1)} = \begin{cases} v_0, & \text{if: } o_{(t)} \text{ in cell } L \leq 0.5 \\ 0, & \text{otherwise.} \end{cases} \tag{3.4a}$$

$$v_{(t+1)} = v_0 \tag{3.4b}$$

#### 4 Clock and three Timers

Initial conditions: $\tau_{1(0)} = 1$, $\phi_{(0)} = 0$, $o_{(0)} = 0$, $\tau_{2(0)} = 0$, $\tau_{3(0)} = 1$, $s_{(0)} = 0$.

$$\tau_{1(t+1)} = \max\left\{\tau_1(t) - h(\tau_{3(t)})(1-W)/T_1,\ 0\right\} \tag{4.1}$$

$$\phi_{(t+1)} = \begin{cases} o_t + \left(\frac{1-e^{-2\tau_{(t)}}}{1-e^{-2}}\right) 1/P \mod 1, & \text{if: } \tau_{1(t)} > 0 \\ 0, & \text{otherwise.} \end{cases} \tag{4.2}$$

$$o_{(t+1)} = \frac{\sin(2\pi(\phi_{(t+1)} - \frac{1}{4})) + 1}{2} \tag{4.3}$$

$$\tau_{2(t+1)} = \begin{cases} 1, & \text{if: } (\tau_{1(t)} > 0) \cap (o_{(t)} > \theta) \\ \max\left\{\tau_{2(t)} - \left(\frac{1-e^{-2\tau_{1(t)}}}{1-e^{-2}}\right) 1/T_2,\ 0\right\}, & \text{if: } (\tau_{1(t)} > 0) \cap (o_{(t)} \le \theta) \\ 0, & \text{otherwise.} \end{cases} \tag{4.4}$$

$$\tau_{3(t+1)} = \begin{cases} \tau_{3(t)}, & \text{if: } \tau_{1(t)} = 0 \\ \max\left\{\tau_{3(t)} - 1/T_3,\ 0\right\}, & \text{otherwise.} \end{cases} \tag{4.5}$$

$$s_{(t+1)} = \begin{cases} s_{(t)}, & \text{if: } s_{(t)} > 0 \\ f(t_{2(t)}), & \text{if: } (s_{(t)} = 0) \cap (\tau_{1(t)} = 0) \\ 0, & \text{otherwise.} \end{cases} \tag{4.6}$$

$$v_{(t+1)} = g(\tau_{3(t)})v_0 \tag{4.7}$$

#### 5 Three Timers and no Clock

Initial conditions: $\tau_{1(0)} = 1$, $o_{(0)} = 0$, $\tau_{2(0)} = 0$, $\tau_{3(0)} = b(x)$, $s_{(0)} = 0$.

$$\tau_{1(t+1)} = \max\{\tau_1(t) - 1/T_1,\ 0\} \tag{5.1}$$

$$o_{(t+1)} = \begin{cases} a(\tau_{3(t)}), & \text{if : } \tau_{1(t)} > 0 \\ 0, & \text{otherwise.} \end{cases} \tag{5.2}$$

$$\tau_{2(t+1)} = \begin{cases} 1, & \text{if: } (\tau_{1(t)} > 0) \cap (o_{(t)} > \theta) \\ \max\left\{\tau_{2(t)} - \left(\frac{1-e^{-2\tau_{1(t)}}}{1-e^{-2}}\right) 1/T_2,\ 0\right\}, & \text{if: } (\tau_{1(t)} > 0) \cap (o_{(t)} \le \theta) \\ 0, & \text{otherwise.} \end{cases} \tag{5.3}$$

$$\tau_{3(t+1)} = \begin{cases} \tau_{3(t)}, & \text{if: } \tau_{1(t)} = 0 \\ \max\left\{\tau_{3(t)} - \left(\frac{1-e^{-2\tau_{1(t)}}}{1-e^{-2}}\right) 1/T_3,\ 0\right\}, & \text{otherwise.} \end{cases} \tag{5.4}$$

$$s_{(t+1)} = \begin{cases} s_{(t)}, & \text{if: } s_{(t)} > 0 \\ f(t_{2(t)}), & \text{if: } (s_{(t)} = 0) \cap (\tau_{1(t)} = 0) \\ 0, & \text{otherwise.} \end{cases} \tag{5.5}$$

#### 6 Model variants

- For the spatial variant of the clock and timer model, substitute (1.1) with:

$$\tau_{1(t+1)} = \begin{cases} 1, & \text{if: } L - x < r \\ (L - x - r)/\lambda, & \text{if: } r < L - x \leq r + \lambda \\ 0, & \text{otherwise.} \end{cases} \tag{6.1}$$

- For a flat timer 2 rate profile (as in main text Figure 4J,L), substitute (2.4) with:

$$\tau_{2(t+1)} = \begin{cases} 1, & \text{if: } (\tau_{1(t)} > 0) \ \cap \ (o_{(t)} > \theta) \\ \max\left\{\tau_{2(t)} - 1/T_2,\ 0\right\}, & \text{if: } (\tau_{1(t)} > 0) \ \cap \ (o_{(t)} \leq \theta) \\ 0, & \text{otherwise.} \end{cases} \tag{6.2}$$

- For the spatial variant of the Clock and three Timers model, substitute (4.1) with:

$$\tau_{1(t+1)} = \begin{cases} 1, & \text{if: } L - x < r \\ (L - x - r)/h(\tau_{3(t)})\lambda, & \text{if: } r < L - x \leq r + g(\tau_{3(t)})\lambda \\ 0, & \text{otherwise.} \end{cases} \tag{6.3}$$

### Simulation details

#### 1 Clock and timer model

| Simulation | $T$ | $\lambda$ | $P$ | $v$ | $n$ | $r$ | $L_0$ |
|---|---|---|---|---|---|---|---|
| Fig 3B; Movie 1 (row 1) | 150 | - | 50 | 0.2 | 2 | 5 | 1 |
| Fig 3C (row 1); Movie 1 (row 2) | 150 | - | 30 | 0.2 | 2 | 5 | 1 |
| Fig 3C (row 2); Movie 1 (row 3) | 250 | - | 50 | 0.2 | 2 | 5 | 1 |
| Fig 3C (row 3); Movie 1 (row 4) | 150 | - | 50 | 0.4 | 2 | 5 | 1 |
| Fig 3D (row 1, left) | 20-500 (20) | - | 50 | 0.2 | 2 | 5 | 1 |
| Fig 3D (row 1, right) | - | 5-100 (5) | 50 | 0.2 | 2 | 5 | 1 |
| Fig 3D (row 2, left) | 20-500 (20) | - | 50 | 0.2 | 2 | 5 | 1 |
| Fig 3D (row 2, right) | - | 5-100 (5) | 50 | 0.2 | 2 | 5 | 1 |
| Fig 3D (row 3, left) | 20-500 (20) | - | 50 | 0.2 | 2 | 5 | 1 |
| Fig 3D (row 3, right) | - | 5-100 (5) | 50 | 0.2 | 2 | 5 | 1 |
| Fig 3D (row 4, left) | 150 | - | 5-100 (5) | 0.2 | 2 | 5 | 1 |
| Fig 3D (row 4, right) | - | 30 | 5-100 (5) | 0.2 | 2 | 5 | 1 |
| Fig 3E (panel 1) | 150 | - | 50 | 0.05-1 (0.05) | 2 | 5 | 1 |
| Fig 3E (panel 2) | - | 75 | 50 | 0.05-1 (0.05) | 2 | 5 | 1 |
| Fig 3E (panel 3) | 150 | - | 50 | 0.05-1 (0.05) | 2 | 5 | 1 |
| Fig 3E (panel 4) | - | 75 | 50 | 0.05-1 (0.05) | 2 | 5 | 1 |
| Movie 2 (row 1) | 100 | - | 50 | 0.2 | 2 | 5 | 1 |
| Movie 2 (row 2) | 100 | - | 50 | 0.5 | 2 | 5 | 1 |
| Movie 2 (row 3) | - | 50 | 50 | 0.2 | 2 | 5 | 1 |
| Movie 2 (row 4) | - | 50 | 50 | 0.5 | 2 | 5 | 1 |

Table 1: **Parameter values for all simulations using the clock and timer model.** $T$ = timer duration (timesteps); $\lambda$ = SAZ lengthscale (cells); $P$ = oscillator period (timesteps); $v$ = elongation rate (timestaps); $n$ = number of switch states; $r$ = posterior signalling centre range (cells); $L_0$ = initial embryo length (cells). Ranges are in the form: first-last (step size).

### 2 Clock and two timers model

| Simulation | $T_1$ | $\lambda$ | $P$ | $\theta$ | $T_2$ | $v$ | $n$ | $r$ | $L_0$ |
|---|---|---|---|---|---|---|---|---|---|
| Fig 4D (row 1); Movie 3 (row 1) | 200 | - | 100 | 0.995 | 95 | 0.2 | 3 | 5 | 1 |
| Fig 4D (row 2); Movie 3 (row 2) | 200 | - | 100 | 0.995 | 95 | 0.2 | 6 | 5 | 1 |
| Fig 4D (row 3); Movie 3 (row 3) | 200 | - | 120 | 0.995 | 95 | 0.2 | 6 | 5 | 1 |
| Fig 4D (row 4); Movie 3 (row 4) | 200 | - | 50 | 0.995 | 95 | 0.2 | 6 | 5 | 1 |
| Suppl Fig 2C,D | 100 | - | 100 | 0.995 | 120 | 0.5 | 3 | 5 | 1 |
| Suppl Fig 2F | 10-250 (10) | - | 100 | 0.995 | 100 | 0.5 | 2 | 5 | 1 |

Table 2: **Parameter values for all simulations using the clock and two timers model.** $T_1$ = timer 1 duration (timesteps); $\lambda$ = SAZ lengthscale (cells); $P$ = oscillator period (timesteps); $\theta$ = oscillator threshold; $T_2$ = timer 2 duration (timesteps); $v$ = elongation rate (timestaps); $n$ = number of switch states; $r$ = posterior signalling centre range (cells); $L_0$ = initial embryo length (cells). Ranges are in the form: first-last (step size).

### 3 Clock and timer model with feedback

| Simulation | $T$ | $\lambda$ | $P$ | $k$ | $v$ | $r$ | $L_0$ |
|---|---|---|---|---|---|---|---|
| Fig 6D; Movie 4 (row 1); Movie 5 (row 1) | 110 | - | 100 | - | 0.25 | 5 | 1 |
| Fig 6E (row ); Movie 4 (row 2) | 110 | - | 100 | - | 0.5 | 5 | 1 |
| Fig 6E (row 2); Fig 6F (left); Movie 4 (row 3) | 190 | - | 100 | - | 0.5 | 5 | 1 |
| Fig 6E (row 3); Movie 4 (row 4) | - | 30 | 100 | - | 0.5 | 5 | 1 |
| Fig 6F (top right) | 2-300 (2) | - | 100 | - | 0.5 | 5 | 1 |
| Fig 6F (bottom right) | - | 2-100 (2) | 100 | - | 0.5 | 5 | 1 |
| Fig 6G (row 1); Fig 6H (top left); Movie 5 (row 2) | 70 | - | 100 | 1.0 | 0.25 | 5 | 1 |
| Fig 6G (row 2); Movie 5 (row 3) | 55 | - | 100 | 2.0 | 0.25 | 5 | 1 |
| Fig 6G (row 3); Movie 5 (row 4) | 85 | - | 100 | 2.0 | 0.25 | 5 | 1 |
| Fig 6H (top right) | 2-300 (2) | - | 100 | 1.0 | 0.25 | 5 | 1 |
| Fig 6H (middle right) | 2-300 (2) | - | 100 | 2.0 | 0.25 | 5 | 1 |
| Fig 6H (bottom) | 70 | - | 100 | 1.0 | 0.25 | 5 | 1 |

Table 3: **Parameter values for all simulations using the clock and timer model with feedback.** $T$ = timer duration (timesteps); $\lambda$ = SAZ lengthscale (cells); $P$ = oscillator period (timesteps); $k$ = feedback strength parameter; $v$ = elongation rate (timestaps); $r$ = posterior signalling centre range (cells); $L_0$ = initial embryo length (cells). Ranges are in the form: first-last (step size).

#### 4 Clock and three timers model

| Simulation | $T_1$ | $\lambda$ | $P$ | $\theta$ | $T_2$ | $T_3$ | $v$ | $n$ | $r$ | $L_0$ |
|---|---|---|---|---|---|---|---|---|---|---|
| Fig 7C (rows 1-3); Movie 6 (row 1) | 200 | - | 100 | 0.995 | 90 | 3000 | 0.25 | 3 | 5 | 1 |
| Fig 7C (row 4); Movie 6 (row 2) | 200 | - | 100 | 0.995 | 90 | 3600 | 0.25 | 3 | 5 | 1 |
| Fig 7C (row 5); Movie 6 (row 3) | 200 | - | 100 | 0.995 | 90 | 2200 | 0.25 | 3 | 5 | 1 |
| Fig 7C (row 6); Movie 6 (row 4) | 200 | - | 100 | 0.995 | 90 | 4400 | 0.25 | 3 | 5 | 1 |
| Fig 7D (left, black); Suppl Fig 3A; Movie 7 (row 1) | 200 | - | 25 | 0.995 | 23 | 1000 | 1 | 6 | 5 | 1 |
| Fig 7D (left, purple); Suppl Fig 3A; Movie 7 (row 2) | - | 200 | 25 | 0.995 | 23 | 1000 | 1 | 6 | 5 | 1 |
| Suppl Fig 3B (black); Movie 8 (row 1) | 200 | - | 25 | 0.995 | 23 | 2000 | 1 | 2 | 5 | 1 |
| Suppl Fig 3B (purple); Movie 8 (row 2) | - | 200 | 25 | 0.995 | 23 | 2000 | 1 | 2 | 5 | 1 |
| Suppl Fig 3C (black); Movie 9 (row 1) | 200 | - | 25 | 0.995 | 23 | 4000 | 1 | 2 | 5 | 1 |
| Suppl Fig 3C (purple); Movie 9 (row 2) | - | 200 | 25 | 0.995 | 23 | 4000 | 1 | 2 | 5 | 1 |
| Fig 7D (right); Movie 10 | 200 | - | 25 | 0.995 | 23 | 2000 | 1 | 2 | 5 | 1 |
| Fig 8D (top); Movie 11 (row 2) | 150 | - | 100 | 0.995 | 95 | 800 | 1 | 6 | 5 | 1 |

Table 4: **Parameter values for all simulations using the clock and three timers model.** $T_1$ = timer 1 duration (timesteps); $\lambda$ = SAZ lengthscale (cells); $P$ = oscillator period (timesteps); $\theta$ = oscillator threshold; $T_2$ = timer 2 duration (timesteps); $T_3$ = timer 3 duration (timesteps); $v$ = elongation rate (timestaps); $n$ = number of switch states; $r$ = posterior signalling centre range (cells); $L_0$ = initial embryo length (cells).

#### 5 Three timers and no clock model

| Simulation | $T_1$ | $\theta$ | $T_2$ | $T_3$ | $n$ | $L_0$ |
|---|---|---|---|---|---|---|
| Fig 8C; Fig 8D (bottom); Movie 11 (top) | 150 | 0.995 | 95 | 800 | 6 | 200 |

Table 5: **Parameter values for simulations using the three timers and no clock model.** $T_1$ = timer 1 duration (timesteps); $\theta$ = oscillator threshold; $T_2$ = timer 2 duration (timesteps); $T_3$ = timer 3 duration (timesteps); $n$ = number of switch states; $L_0$ = initial embryo length (cells).
