## Supplementary Figures for "Time and Space in Segmentation"

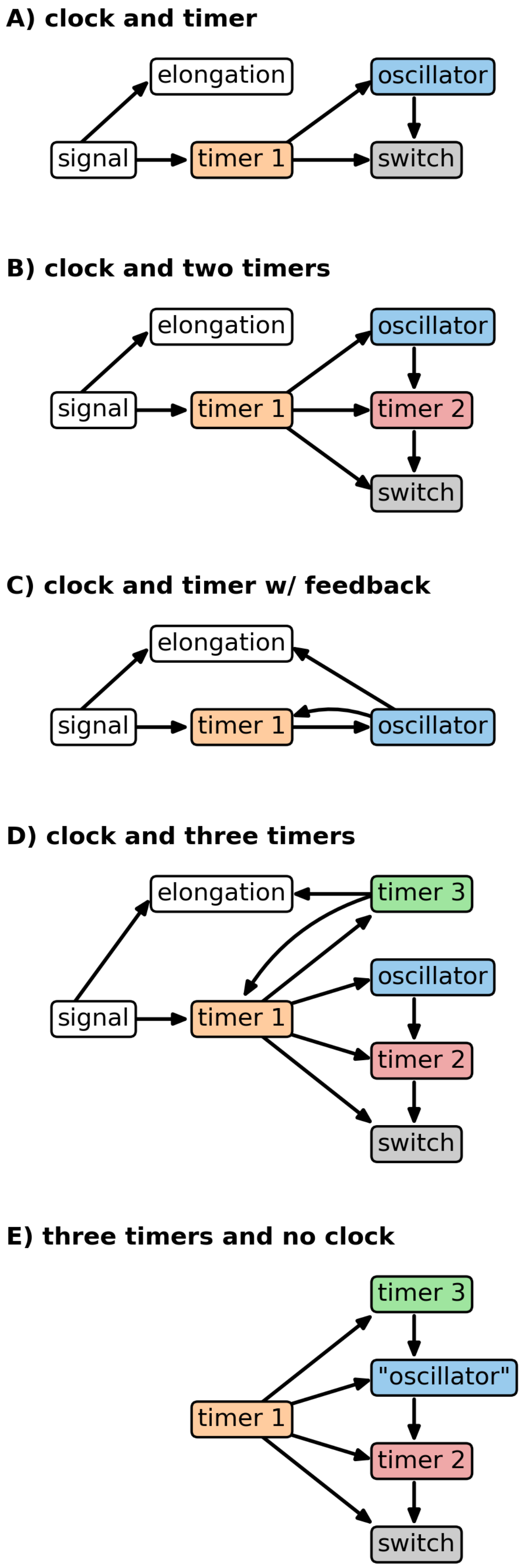


Supplementary Figure 1: **Summary diagrams of all models.** Dynamical modules are shown as coloured boxes, other model features are shown as white boxes. Regulatory connections are indicated with arrows.

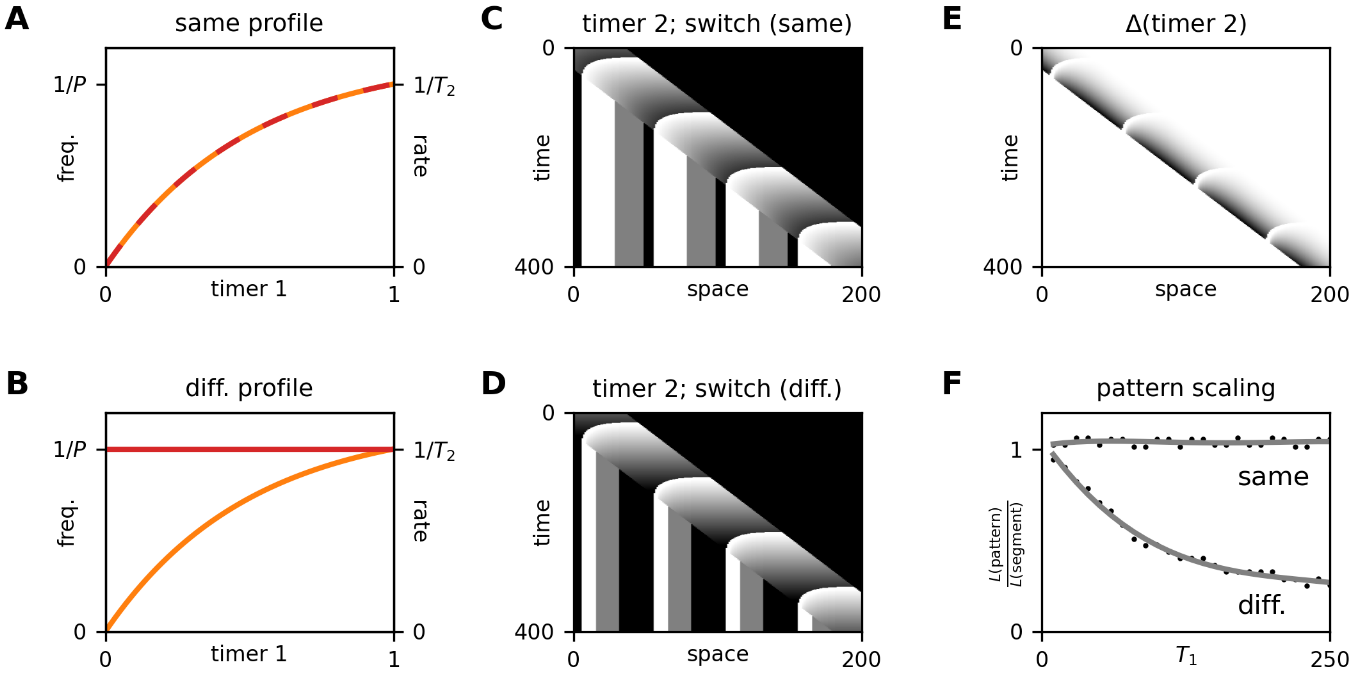


Supplementary Figure 2: **Additional results from the clock and two timers model.** **A,B:** rate profile of timer 2 (red) compared to frequency profile of the oscillator (orange). In A, the module dynamics are affected identically by timer 1; in B, they are affected differently. **C,D:** kymographs showing the output from timer 2 (SAZ pulses) and the switch (stable stripes), using either the profiles from A (C) or from B (D). White=1; black=0. In both simulations, $T_2$ is moderately larger than $P$, and the timer 2 : switch mapping consists of three equally-sized states. The different timer 2 rate profiles result in different segment polarity patterns: in C, lengthscale of the pattern is longer than the repeat length, and so the posterior state (black) is truncated; in D, the lengthscale of the pattern is shorter than the repeat length, and so the posterior state is expanded. **E:** kymograph showing the magnitude of the difference between the timer 2 states in C and the timer 2 states in D. Darker shade indicates greater difference. Note that the states diverge towards the anterior of the SAZ, because the timer 2 in D ticks faster than the timer 2 in C. **F:** plots showing the ratio between the lengthscale of the segment-polarity pattern and the length of the segment repeat, for different value of $T_1$ (timer 1 duration). If the rate and frequency profiles are the same ("same", see panel A), the relationship is flat (*i.e.*, $T_1$ has no effect on the pattern). If the rate and frequency profiles are different ("diff.", see panel B), the ratio decreases with larger values of $T_1$, because a larger $T_1$ means a longer time in which timer 2 is ticking faster than the oscillator. Black dots show individual simulations; grey lines show fitted polynomials. Deviations are due to the discrete nature of the simulation output.

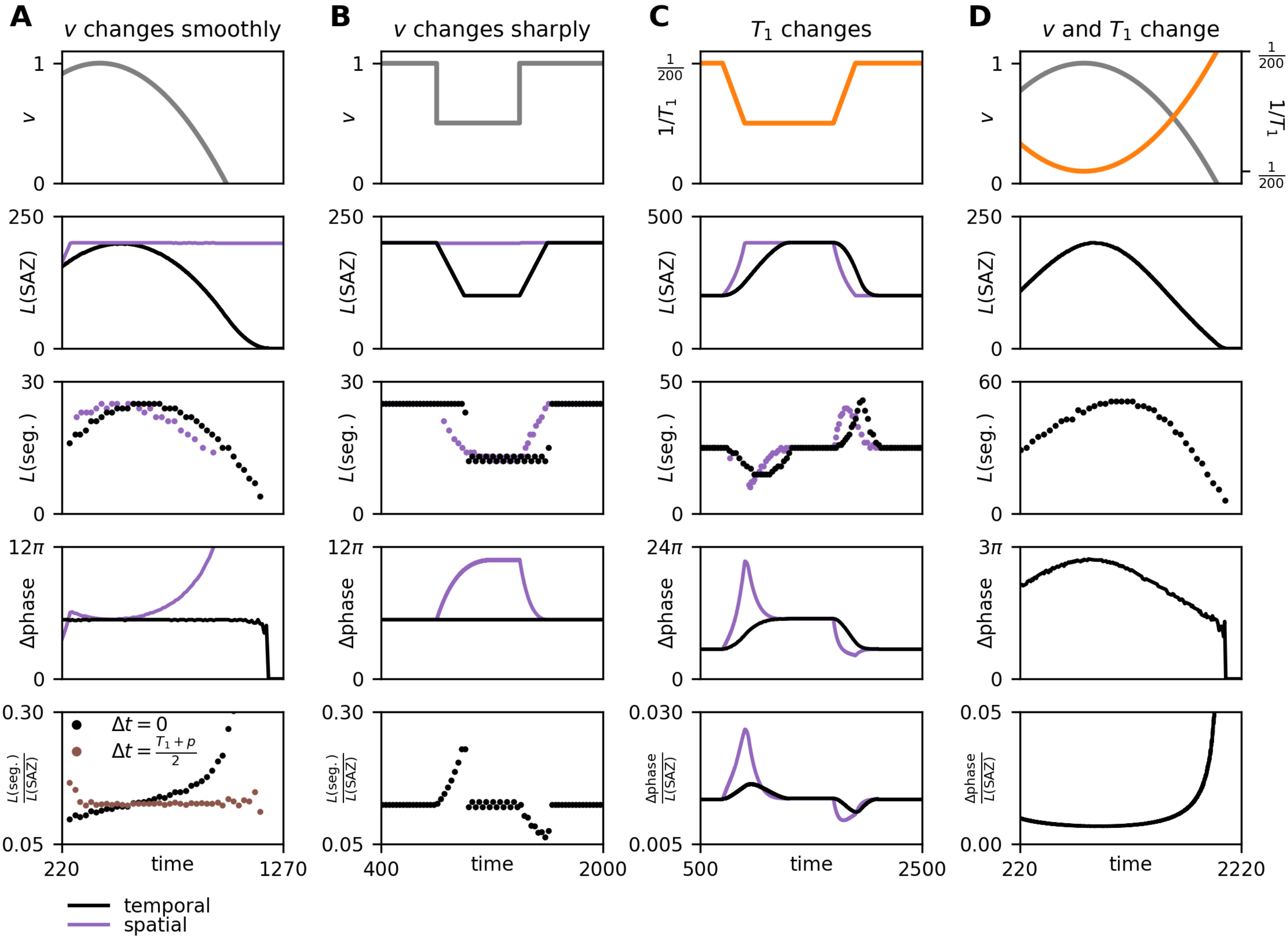


Supplementary Figure 3: **Additional results from the clock and three timers model.** **A:** summary statistics over time for a simulation (see Movie 7) where the elongation rate (grey line) changes smoothly over time. With the temporal model variant (black line), SAZ size mirrors the elongation rate, the SAZ maintains a constant phase profile, and the SAZ shrinks and disappears after elongation halts. With the spatial variant (purple line), the SAZ size remains constant, the SAZ phase profile varies dramatically, and the SAZ is maintained after elongation halts, continuously accumulating additional stripes. For both variants, the segment length profile mirrors the elongation rate profile, but there is a larger time lag for the temporal variant. With the temporal variant, segment length varies as a proportion of SAZ length over time (bottom panel). However, the changing relationship is largely due to the time delays involved in segment patterning: if segment length is plotted as a fraction of the SAZ size $(T_1 + P)/2$ timepoints earlier (brown dots), the relationship is approximately constant. **B:** summary statistics over time for a simulation (see Movie 8) where the elongation rate changes sharply. With the temporal variant, there is a long delay (determined by $T_1$) before an elongation rate perturbation affects segment size, and the change, when it eventually occurs, is sudden. With the spatial variant, segment size is affected immediately, and there is a smooth segment size adjustment over time. **C:** summary statistics over time for a simulation (see Movie 9) where the timer 1 decrease rate (or $1/\lambda$ value, for the spatial model variant) changes as shown (orange line). For both temporal and spatial model variants, segment size is independent of $T_1$ (or $\lambda$) at "steady state", but there are transient effects on segment size while the system adjusts to a change. With the temporal variant, this is because the wavefront temporarily moves faster or slower than the elongation rate while the size of the SAZ is changing. With the spatial variant it is because the frequency profile becomes transiently perturbed. With both models, the new "steady state" is associated with a new SAZ size and frequency profile. **D:** summary statistics over time for a simulation (see Movie 10) combining both elongation rate changes (grey line) and timer 1 decrease rate changes (orange line). Curved profiles of SAZ length, segment length, and SAZ phase difference all result.
